# Multiscale imaging and quantitative analysis of plasma membrane protein-cortical actin interplay

**DOI:** 10.1101/2023.01.22.525112

**Authors:** Aparajita Dasgupta, Huong-Tra Ngo, Deryl Tschoerner, Nicolas Touret, Bruno da Rocha-Azevedo, Khuloud Jaqaman

**Affiliations:** Department of Biophysics, University of Texas Southwestern Medical Center; Dallas, TX, USA; Department of Biochemistry, University of Alberta; Edmonton, AB, Canada; Lyda Hill Department of Bioinformatics, University of Texas Southwestern Medical Center; Dallas, TX, USA

## Abstract

The spatiotemporal organization of cell surface receptors is important for cell signaling. Cortical actin (CA), the subset of the actin cytoskeleton subjacent to the plasma membrane (PM), plays a large role in cell surface receptor organization. This was however shown largely through actin perturbation experiments, which raise concerns of nonspecific effects and preclude quantification of actin architecture and dynamics under unperturbed conditions. These limitations make it challenging to predict how changes in CA properties can affect receptor organization. To derive direct relationships between the architecture and dynamics of CA and the spatiotemporal organization of PM proteins, including cell surface receptors, we developed a multiscale imaging and computational analysis framework based on the integration of single-molecule imaging (SMI) of PM proteins and fluorescent speckle microscopy (FSM) of CA (combined: SMI-FSM) in the same live cell. SMI-FSM revealed differential relationships between PM proteins and CA based on the PM proteins’ actin binding ability, diffusion type and local CA density. It also highlighted the complexity of cell wide actin perturbation, where we found that global changes in actin properties caused by perturbation were not necessarily reflected in the CA properties near PM proteins, and the changes in PM protein properties upon perturbation varied based on the local CA environment. Given the widespread use of SMI as a method to study the spatiotemporal organization of PM proteins and the versatility of SMI-FSM, we expect it to be widely applicable to enable future investigation of the influence of CA architecture and dynamics on different PM proteins, especially in the context of actin-dependent cellular processes, such as cell migration.

**Significance:** Plasma membrane protein organization, an important factor for shaping cellular behaviors, is influenced by cortical actin, the subset of the actin cytoskeleton near the plasma membrane. Yet it is challenging to directly and quantitatively probe this influence. Here, we developed an imaging and analysis approach that combines single-molecule imaging, fluorescent speckle microscopy and computational statistical analysis to characterize and correlate the spatiotemporal organization of plasma membrane proteins and cortical actin. Our approach revealed different relationships between different proteins and cortical actin, and highlighted the complexity of interpreting cell wide actin perturbation experiments. We expect this approach to be widely used to study the influence of cortical actin on different plasma membrane components, especially in actin-dependent processes.

## Introduction

The spatiotemporal organization of cell surface receptors plays an important role in cell signaling (*1-4*). It modulates ligand binding (*5, 6*), homotypic and heterotypic receptor interactions (*7-9*), and recruitment of downstream effectors (*1, 3, 10*). Multiple factors affect cell surface receptor organization and function, including interactions with other plasma membrane (PM) proteins and lipids (*11-13*), the extracellular matrix (*14, 15*) and the cortical cytoskeleton (*16-18*). Cortical actin (CA), a major constituent of the cortex, exerts both direct influence on PM proteins, and indirect influence through its effect on lipids (*19-21*). Its influence is in part described by the picket-fence model, where the filamentous CA meshwork acts as a barrier to the diffusion of PM proteins and lipids (*15, 22-24*). However, the cortex is a dynamic structure that consists of more than a mere meshwork; CA filaments also form dynamic structures that may influence PM protein (and lipid) movement and clustering (*25-29*). At the same time, cell surface receptor signaling can regulate CA remodeling (*30, 31*). Thus, studying the dynamic organization of and interplay between CA and PM proteins is critical for a comprehensive understanding of cell surface receptor signaling.

A limiting factor for studying the interplay between CA and the spatiotemporal organization of the PM is that the CA meshwork is too dense to simultaneously resolve its architecture and dynamics with conventional light microscopy (*32*). Therefore, insights into the role of the cortex in PM organization have come largely from perturbation-based studies (*22, 29, 33*). Single-molecule (SM) imaging (SMI) and tracking of several PM molecules have demonstrated that perturbing actin leads to faster diffusion of these molecules, or an increase in their freely moving population, in line with CA acting as a barrier (*34, 35*). However, perturbations preclude the quantification of CA architecture and dynamics under unperturbed conditions. They also compromise PM integrity and cell health.

To complement/circumvent perturbation experiments, a few previous studies employed simultaneous imaging of PM molecules and CA to directly visualize the correlation between CA and the spatiotemporal organization of PM molecules. By combining SMI of PM proteins with conventional imaging of actin, it was observed that B cell receptors were less mobile in actin-rich regions vs. in actin-poor regions (*34*) or that FCεRI receptors were excluded from crossing into actin-rich regions (*36*). To probe CA at higher resolution, one study employed a combination of super-resolution radial fluctuations (SRRF), fluorescence correlation spectroscopy and number and brightness analysis (*37*). However, while actin disruption by latrunculin A (Lat A) altered the mobility of EGFR in this study, the employed analyses found no direct correlation between the diffusion or clustering of EGFR and CA as imaged by SRRF. As such, the influence of actin on EGFR mobility was inferred to be indirect, through the influence of actin on the PM (*37*). Meanwhile, Li et al. (*24*) used fluorescent magnetic nanoparticles to drag and image PM proteins across the PM, followed by super-resolution imaging of CA using dSTORM. They found that PM protein movement encountered resistance in areas with dense CA filaments. However, despite the co-imaging of PM proteins and CA, these studies provided mostly a static view of CA architecture, thus limiting the characterization of CA dynamics.

To observe the spatiotemporal organization of PM proteins and, simultaneously, the architecture and dynamics of CA with high spatiotemporal resolution, we combined SMI of PM proteins with quantitative fluorescent speckle microscopy (qFSM) of CA in individual live cells. In qFSM, actin monomers are labeled sub-stoichiometrically, so that speckles are formed when several monomers come together due to actin polymerization. QFSM allows quantitative characterization of CA architecture, turnover and movement (*38*), and is thus well-suited to probe CA dynamics alongside the spatiotemporal organization of PM proteins. We then developed a computational analysis pipeline to parse the acquired data, explore the multidimensional dataset of dynamic PM protein properties and associated CA properties, and derive quantitative relationships between them. We applied this framework, collectively called SMI-FSM, to model transmembrane proteins with known actin binding properties, and to the receptor CD36 (*39*), the cell surface mobility, clustering and signaling of which, in multiple cell types, have been shown to depend on actin (*5, 7*). These prior studies however employed actin perturbation, leaving open questions about the direct relationship between CA and CD36 behavior at the PM, which SMI-FSM is ideally suited to address.

## Materials and Methods

### Plasmids

Plasmid encoding CD36-fused at the N-terminus to HaloTag (Halo-CD36) was generated by PCR-cloning the HaloTag coding sequence (primers details in **Table S1**, Integrated DNA Technologies) from pFN21K HaloTag® CMV Flexi® Vector (Promega, Madison, WI), followed by ligation in place of the mApple sequence in the mApple-CD36-C-10 vector (Addgene, Watertown, MA plasmid # 54874 (*5*)), using restriction enzymes AgeI and BglII (New England Biolabs, Ipswich, MA). To generate Halo-TM-ABD and Halo-TM-ABD*, Halo-CD36 plasmid was linearized using restriction enzymes EcoRI and BglII (New England Biolabs) to release the CD36 fragment. TM-ABD or TM-ABD* fragments were then seamlessly cloned (Takara bio, San Jose, CA) into the linearized HaloTag vector by PCR amplification (primers details in **Table S1**, Integrated DNA Technologies) from GFPTM-Ez-AFBD (ABD) and GFPTM-Ez-AFBD* (ABD*) with corresponding flanking homology. GFPTM-Ez-AFBD and GFPTM-Ez-AFBD* were a generous gift from Satyajit Mayor Lab (NCBS, India) (*25*). PLVXCMV100mNeonGreenActin, pSPAX2 and pMD2G plasmids were a generous gift from Gaudenz Danuser lab (The University of Texas Southwestern Medical Center, Dallas, Texas) (*40*).

### Cell lines and cell culture

Human Telomerase (hTERT)-immortalized microvascular endothelial cells isolated from human foreskin (TIME cells, ATCC, Manassas, VA), or TIME cells stably expressing mNeonGreen-Actin (TIME-mNGrActin; see description of cell line generation below), both at passage 16 or higher (unless indicated otherwise), were grown in ATCC’s vascular cell basal medium supplemented with microvascular endothelial cell growth kit-VEGF and 12.5 μg/mL blasticidine (Sigma-Aldrich, St. Louis, MO, 203351) for 48 h at 37°C + 5% CO_2_ until reaching 80-90% confluency. Then cells were passaged and plated on fibronectin (FN)-coated (10 µg/ml, MilliporeSigma, Burlington, MA), base/acid cleaned, 0.17 mm (#1.5) glass bottom dishes (14 mm glass diameter, MatTek, Ashland, MA) or 6 well plates for 18 h prior to experiments.

**Generation of TIME-mNGrActin cell line.** Lentiviral transduction was performed to generate TIME cells stably expressing mNeonGreen tagged actin (TIME-mNGrActin). Briefly, HEK293T cells (Danuser lab) were transfected using 3 µl/µg Polyethylenimine (PEI) (Sigma Aldrich, 408727) with PLVX-mNeonGreenActin-CMV-100 (4 µg) along with VSV envelope vector pSPAX2 (2 µg) and viral packaging construct pMD2G (2 µg). PLVX-mNeonGreenActin-CMV-100 has the fluorescent fusion protein expressed from a truncated CMV promoter containing only the first 100 base pairs from the 5’ end of the CMV promoter (*40*). Viral media were harvested after 48 h, filtered (0.45 µm pore size) and mixed with 0.5 µl/ml dilution of Polybrene (MilliporeSigma, TR-1003-G). To generate the TIME-mNGrActin cell line, TIME cells at 50% confluency were incubated with viral media mixed with polybrene for 72 h. Infected cells at the lower range of mNeonGreen-actin expression were enriched using fluorescence activated cell sorting using a FACSAria system (Children’s Research Institute Flow Cytometry Facility, The University of Texas Southwestern Medical Center, Dallas, TX).

### Transient transfection

For experiments comparing CD36, TM-ABD, and TM-ABD*, TIME-mNGrActin cells (1×10^5^ cells), seeded for 18 h on fibronectin-coated glass bottom dishes, were transiently transfected with Halo-CD36 (0.25 µg), Halo-TM-ABD (0.25 µg) or Halo-TM-ABD* (0.25 µg) using Transfex transfection reagent (ATCC, ACS-4005). Experiments were performed 2 days after transfection. For experiments involving Lat A treatment and the corresponding untreated cells, TIME-mNGrActin cells at 80-90% confluency were transiently transfected in suspension with Halo-CD36 (1 µg) using Universal transfection reagent (Sigma Aldrich, T0956). Transfected cells were passaged the next day and 1×10^5^ cells were seeded on fibronectin-coated glass bottomed dishes. Experiments were performed 2 to 3 days after transfection.

### Halo labeling for SMI

Halo-CD36, Halo-TM-ABD, or Halo-TM-ABD* transfected TIME-mNGrActin cells plated on fibronectin-coated glass bottom dishes for 20-24 h were incubated with 3 nM JF549-Halo ligand (experiments comparing CD36, TM-ABD and TM-ABD*) or 15 nM JF549-Halo ligand (experiments involving Lat A treatment and corresponding untreated cells) (JF549-Halo was a generous gift from Dr. Luke Lavis, Janelia Research Campus, Ashburn, VA) (*41*) for 15 min in complete culture medium. This was followed by 3 quick washes in sterile DPBS. Cells were then incubated for 15 min in dye free complete culture medium. Incubations were at 37°C + 5% CO_2_. Finally, cells were washed 3 times (5 min each) with wash buffer (HBSS + 1 mM HEPES and 0.1% NGS (Normal Goat Serum) for live cells). Cells were incubated in imaging buffer (Oxyfluor 1%, Glucose 0.45% and Trolox 2nM) to reduce photobleaching before and during imaging.

### Drug treatments and perturbations

For cell fixation with paraformaldehyde (PFA), cells were incubated with a 4% PFA solution made in PBS (Electron Microscopy Sciences, 15710, Hatfield, PA) for 15 min at room temperature and then washed three times (5 min each) with wash buffer (HBSS + 0.1% NGS for fixed cells) before adding imaging buffer. For cell fixation with PFA+glutaraldehyde, cells were incubated with a 4% PFA + 0.1% glutaraldehyde (Electron Microscopy Sciences, 16020) solution made in PBS, for 20 min at 4°C, in the dark. Freshly prepared quenching solution (0.1% NaBH4, Oakwood Chemical, 042896, Estill, SC) was added immediately after fixation and cells were incubated for 7 min. Samples were then washed three times (5 min each) at RT with wash buffer. For Lat A treatment, TIME-mNGrActin cells expressing Halo-CD36 labelled with JF549-Halo ligand were incubated with 30 nM Lat A (ThermoFisher Scientific, Waltham, MA, L12370), diluted in imaging buffer, at the microscope, as described in the following section.

### Simultaneous SMI-FSM data acquisition

TIME-mNGrActin cells expressing Halo-CD36, Halo-TM-ABD, or Halo-TM-ABD* and labelled with Halo ligand were imaged at 37°C using an S-TIRF system (Spectral Applied Research, via BioVision Technologies, Exton, PA) mounted on a Nikon Ti-E inverted microscope with Perfect Focus and a 60x/1.49 NA oil TIRF (total internal reflection fluorescence) objective (Nikon Instruments, Melville, NY). The system was equipped with two Evolve EMCCD cameras (Photometrics, Tucson, AZ) for one-color or simultaneous two-color imaging, registered to within 1-3 pixels of each other. A custom 3x tube lens was employed to achieve an 89 nm x 89 nm pixel size in the recorded image. Illumination by a 561 nm diode pumped solid state laser (Cobolt) and/or a 488 nm diode laser (Coherent) was achieved through an ILE laser merge module (Spectral Applied Research, Ontario, Canada), with 561 nm and 488 nm laser power of 13.2 mW and 3.65 mW, respectively, at the coverslip. The penetration depth was set to 80 nm via the Diskovery platform control (Spectral Applied Research). Videos were acquired in live stream mode with MetaMorph (Molecular Devices, San Jose, CA). The 561 channel was used for acquiring SMI videos at 10 Hz for 50 s (i.e. 501 frames) or 90 s (i.e. 901 frames). Simultaneously, the 488 channel was used for acquiring FSM videos at 0.2 Hz for the corresponding 11 or 19 frames. All videos were acquired using an EM gain of 100. Temperature and humidity were maintained using an environment chamber (Okolab, Otaviano, Italy), maintaining cell viability for the duration of the experiments. Single bandpass emission filters, ET520/40 and ET605/52 (Chroma Technology, Bellows Falls, VT), and a dichroic (565dcxr, Chroma Technology) to direct the emission beam to the two cameras, were used in the emission path.

For the CD36, TM-ABD and TM-ABD* datasets, multiple cells were acquired sequentially (for 50 s each) over approximately 40 min of imaging. For the Lat A and its corresponding unperturbed datasets, a single cell was selected from the dish and followed over time, where it was imaged for 3 instances of 1.5 min each, with an ∼5 min resting interval between the imaging instances. The total imaging session in this case, including actual imaging and resting intervals, was about 25 min. In the Lat A dataset, Lat A was added to the sample at the microscope, after selecting a cell for imaging. The first imaging instance started almost immediately after Lat A addition. The time at which Lat A was added was recorded and used to calculate the time post perturbation.

Every SMI-FSM movie was preceded by a brightfield snapshot of the imaged cell region to visually check cell viability.

### FSM of actin with and without fixation

TIME-mNGrActin cells were either left untreated or fixed using PFA + glutaraldehyde (as described above). Both live and fixed cells were then imaged at 37°C using an Olympus IX83 TIRF microscope equipped with a Z-Drift Compensator and a UAPO 100x/1.49 NA oil-immersion TIRF objective (Olympus, Tokyo, Japan). The microscope was equipped with an iXon 888 1k × 1k EMCCD Camera (Andor; Oxford Instruments). With an additional 1.6x magnification in place, the pixel size in the recorded image was 81 nm x 81 nm. Using the Olympus CellSens software, excitation light of 491 nm from an Olympus CellTIRF-4Line laser system was directed to the sample by a TRF8001-OL3 Quad-band dichroic mirror. The penetration depth was set to 70 nm via the cellSens software (Olympus). Laser power was 2.31 mW at the coverslip. FSM videos were acquired at a frame rate of 0.2 Hz for 109 frames (9 min). This was equivalent to the FSM arm of the SMI-FSM data acquisition scheme described above, although with longer acquisition time. Fluorescence was collected, filtered with an emission filter of ET520/40m (Chroma Technology) and projected onto a section of the camera chip by an OptoSplit III 3-channel image splitter (Cairn Research, Faversham, UK). Camera EM gain was set to 100 for all acquisitions.

### Actin FSM + phalloidin imaging

TIME-mNGrActin cells plated on fibronectin coated glass bottomed dishes were fixed with 4% PFA (as described above). Following fixation, sample was incubated with Alexa Fluor™ 647 phalloidin (ThermoFisher Scientific, A22287) at 1:20 dilution in blocking buffer (1% BSA, 5% NGS in wash buffer). After washing, cells were incubated in imaging buffer and imaged at 37°C using an Olympus IX83 TIRF microscope (microscope described in previous section). For excitation, light of 640 nm (0.2 mW at coverslip) and 491 nm (2.3 or 5.1 mW at coverslip) from an Olympus CellTIRF-4Line laser system was directed to the sample. The penetration depth was set to 100 nm via the cellSens software (Olympus). First, the 640 channel was imaged for an exposure time of 99 ms. Then, the 491 channel was used for acquiring FSM videos at 10 Hz for 5 frames. Fluorescence of each wavelength was collected, filtered with emission filters of ET520/40m and ET705/72m (Chroma Technology), and projected onto different sections of the camera chip by an OptoSplit III 3-channel image splitter (Cairn Research). Camera EM gain was set to 100 for all acquisitions.

### Immunofluorescence (IF) imaging of endogenous CD36 and VEGFR2 expression

TIME cells plated for 20 h were washed once in wash buffer to remove the culture medium and then fixed with a 4% PFA solution made in PBS for 15 min at RT. After three washes, samples were blocked for 15 min in blocking buffer and then incubated with primary antibodies, either against CD36 (1:400, FA6-152, Abcam, Cambridge, UK, ab17044) or VEGFR2 (1:200, ab9530, Abcam) for 1h at RT. After three washes, samples were incubated with Alexa Fluor 488-conjugated secondary antibodies (1:1000, ThermoFisher Scientific) for 15 min at RT. Finally, after three subsequent washes, dishes were incubated with imaging buffer to reduce photobleaching. IF imaging was then performed using the S-TIRF microscope described above (section “**Simultaneous SMI-FSM data acquisition**”) using a 488 nm laser power of 2.6 mW at the coverslip.

### IF imaging and analysis of phospho-Src enrichment at CD36 spots

TIME cells, transiently transfected with Halo-CD36 for 2 days, were serum-starved for 3 h. After 2 h of starvation, cells were labelled with 10 nM JF549-Halo ligand (as described in the section “**Halo labeling for SMI**”). The samples were then exposed to either 10 nM thrombospondin-1 (TSP-1) (3074-TH, R&D Systems) or HBSS for 10 min at 37°C, for “TSP-1” or “Control” conditions, respectively. After 10 min, the samples were washed with wash buffer and then fixed with a 4% PFA + 0.1% glutaraldehyde solution made in PBS, for 20 min at 4°C, in the dark. Freshly prepared quenching solution was added immediately after fixation and cells were incubated for 7 min. Samples were then washed three times (5 min each) at RT with wash buffer. Samples were permeabilized using 0.1% Triton X-100. After 3 washes with wash buffer, samples were blocked for 15 min in blocking buffer and then incubated with Alexa Fluor conjugated primary antibody against phospho-Src (Tyr419) (44-660G, ThermoFisher Scientific). Finally, after three subsequent washes, dishes were incubated with imaging buffer to reduce photobleaching. IF imaging was performed as describe above, using laser powers of 13.2 mW for the 561 channel (CD36) and 3.65 mW for the 488 channel (pSrc) at the coverslip. The two channels were imaged sequentially (561 followed by 488).

For the analysis of pSrc enrichment at CD36 spots from the acquired IF images, the centers of CD36 spots were determined using “point-source detection” in u-track (*42, 43*) (https://github.com/DanuserLab/u-track). In the detection step, default parameter values were used, except for the α-value of 0.01 for determining object detection significance by comparing the fitted amplitude to the local background noise distribution. A PSF sigma of 1.228 pixels was used. Enrichment analysis was carried out as described in the “Intensity enrichment analysis” section of (*5*). To compare the control and TSP-1 stimulated datasets to each other and to their randomizations, multiway analysis of variance (ANOVA) with default linear model was performed (MATLAB function anovan()) followed by Tukey-Kramer post hoc analysis.

### SM particle detection and tracking

SM tracks were constructed from the SMI streams using u-track (*42*) (Version 2.2.1; https://github.com/DanuserLab/u-track). Detection was achieved using the “gaussian mixture-model fitting” option. The SM detection and tracking parameters were optimized based on the performance diagnostics included in u-track and visual inspection of the results. The mean SM localization precision, as estimated during the Gaussian mixture-model fitting step (*42*), was 16-24 nm. **Tables S2** and **S3** list, respectively, the detection and tracking parameters with non-default values.

### SM particle track refinement and processing

The SM tracks were processed as follows in preparation for integrating the SM tracks with speckle information:

***Step 1.*** Tracks with duration < 10 timepoints were discarded. These tracks were too short to be informative in the ensuing analyses.

***Step 2.*** Artifactual merging and splitting events between tracks were eliminated. This included the occasional instances of a merge and a split occurring with the same track at the same time, and the following artifacts (as described previously (*44*)): (i) a track appearing and then merging with another track in the very next frame, (ii) a track splitting from another track and then re-merging with it in the very next frame, and (iii) a track splitting from another track and then disappearing immediately.

***Step 3.*** SM apparent assembly state was estimated for each track over time. The apparent assembly state of each SM track was calculated from its particle intensities and sequence of merging and splitting events, as described previously (*44, 45*). The assembly state was ‘‘apparent’’ because, by definition, only labeled molecules were visualized and counted. In brief, this analysis utilized the mean intensity of an individual fluorophore, which was estimated per SMI movie by decomposing the distribution of detected SM particle intensities into multiple modes corresponding to 1, 2, 3, etc. fluorophores. Each mode was taken as a log-normal distribution, where the mean and standard deviation of mode n were (approximately) n times those of mode 1. The fit was achieved using least-squares, and the number of modes was determined using the Bayesian Information Criterion. With this, the mean individual fluorophore intensity was obtained per SMI movie, and then, together with the tracks’ merging and splitting history, it was used to estimate the apparent assembly state of each SM track (*44*).

***Step 4.*** Tracks were segmented at merging and splitting events. Molecular interactions (as reflected by merging and splitting events) may change molecular behavior, and possibly the crosstalk between the molecules and the CA. Thus, after estimating the apparent assembly state of the SM tracks, the SM tracks were segmented at merging and splitting events, so that from now on each SM track had one, constant assembly state. For example, if two SMs merged and then stayed together for the rest of the movie, they would provide three SM tracks, one for each molecule before the merging event, and one for the joint particle after the merging event. Note that the average assembly state over all SM tracks was close to 1, confirming the SM nature of our labeling and imaging scheme (Fig. S1A).

***Step 5.*** Tracks outside of the cell region of interest (ROI) mask were discarded. For SMI-FSM data integration and analysis, only SM tracks inside the cell ROI mask (determined from the images in the actin speckle channel) were utilized. An SM track was taken as inside the cell ROI mask if its center position was inside the cell ROI mask.

### QFSM analysis

FSM time-lapse sequences were analyzed using the qFSM software package (*38*), following the instructions in (*46*). Briefly, first noise model calibration was performed to estimate spatial correlation in background noise (*46*) (Step 1). Then cell masks were generated either within the qFSM package at the thresholding and mask refinement steps (Steps 2 and 3) or they were hand-drawn (if thresholding failed). Then speckles were detected (Step 4) and tracked (Step 5) within the cell mask. The flow tracking step was not utilized as the speckles in our movies did not exhibit collective flow (Step 6). Polymerization/depolymerization events were tracked to generate kinetic maps (Step 7) (not used for further analysis). The qFSM parameters for the relevant analysis steps, were optimized based on visual inspection of the results. Parameters with non-default values are listed in **Tables S4-S6**.

### Analysis of individual actin speckle properties derived from qFSM

This analysis started with the speckle tracks as obtained from qFSM package (*38, 46*). After removing “ghost speckles,” i.e. speckles that existed for one frame only (*47*), the following properties were calculated for each speckle:

- Position (x- and y-coordinates) per FSM frame.
- Intensity per FSM frame.
- Local background intensity per FSM frame.
- Frame-to-frame displacement per FSM frame. For FSM frame i, the displacement was defined as the change in position from FSM frame i to FSM frame i+1. A speckle had no displacement information at its last frame. As a quality control for speckle frame-to-frame displacement, we estimated the apparent speckle frame-to-frame displacement resulting from speckle localization and tracking errors, as measured in cells fixed with PFA+glutaraldehyde. We found that ∼75% of speckles in fixed cells exhibited frame-to-frame displacements of magnitude ≤ 2^1/2^ pixels (Fig. S1B). Thus, to remove artifactual speckle movement, individual speckle frame-to-frame displacements with magnitude < 1.5 pixels were set to 0 (*48*) (this amounted to ∼40% of individual speckle displacements in live cells; Fig. S1B).
- Speckle lifetime. This was calculated as the number of FSM frames that a speckle track existed (this property was for the whole track, and not per frame). For speckles that appeared after the first FSM frame and disappeared before the last FSM frame, the calculated lifetime was their true, complete lifetime. But for speckles that existed in the first and/or last frame, the calculated lifetime was only a minimum estimate of their lifetime, i.e., their lifetime information was incomplete. Because only 33-47% of speckles had complete lifetimes in our movies, we eventually did not employ the speckle lifetime as a speckle property beyond some initial analyses.

### Calculation of global actin speckle properties

Using the individual actin speckle properties, the following global actin speckle properties were calculated. These properties are referred to as global because they describe the speckles throughout the cell ROI mask regardless of the vicinity of any SM molecule.

- Global speckle displacement magnitude. This was viewed as either the mean of all individual speckle displacement magnitudes per cell (over all frames), or as the distribution of individual speckle displacement magnitudes over all cells.
- Global speckle intensity. This was viewed as the mean of all individual speckle intensities per cell (over all frames).
- Global speckle density. For this, first the speckle density was calculated for each frame of a cell as the number of speckles within the cell mask divided by the cell mask area in that frame. Then, the global speckle density for the cell was taken as the mean speckle density over all frames.

### Single-cell multivariate regression (MVRG) analysis of SM tracklet and local speckle properties

Linear MVRG (using the MATLAB function mvregress()) was performed on an individual cell (movie) basis. SM tracklet properties were taken as the response (“dependent”) variables and the associated local speckle properties were taken as the design matrix (“independent variables”). Each property was normalized by its sample mean and standard deviation before MVRG analysis. For any combination of SM tracklet and speckle properties, only complete datapoints (i.e., datapoints without any missing values for any of the properties of interest) were utilized. MVRG was performed only when there was a minimum of 11 complete datapoints. To check multicollinearity in the design matrices (i.e. interdependence between the “independent variables”), the variance inflation factor was calculated for each performed MVRG (*49*).

MVRG yielded per-single cell coefficients quantifying the extent of (mathematical) dependence of any SM tracklet property on the co-analyzed local speckle properties (one coefficient per SM tracklet-actin speckle property pair). The MVRG coefficients of the set of cells representing a particular experimental condition were then grouped for subsequent analysis of their significance and their comparison between experimental conditions. Upon grouping, outlier single-cell MVRG coefficients were identified using the generalized extreme Studentized deviate test, useful for detecting multiple outliers in a univariate dataset that follows an approximately normal distribution (*50*) (MATLAB function isoutlier(data, ‘gesd’)).

### Statistical tests for comparing SM or speckle properties between different datasets

To compare the SM or speckle properties between two experimental conditions, a two-sample t-test was performed (MATLAB function ttest2()). If more than two experimental conditions were compared, multiway analysis of variance (ANOVA) with default linear model was performed (MATLAB function anovan()), followed by Tukey-Kramer post hoc analysis (MATLAB function multcompare()).

To compare the SM or speckle properties between the “Free” and “Confined” subsets of SM tracklets of an experimental condition, a paired-sample t-test was performed (MATLAB function ttest()). A paired-sample t-test was appropriate in this case as each movie provided measurements for the “Free” subset and the “Confined” subset.

For SM and speckle properties where multiple instances of the same cell were imaged, to compare between two experimental conditions at a particular instance, a two-sample t-test was performed. To compare between two experimental conditions across all instances, the mean over all cells at each instance was calculated per condition, and then the two groups of per-instance means were compared using a paired-t-test (paired by instance).

For all of the above tests, comparisons with a p-value ≤ 0.05, 0.01 or 0.001 were indicated in the figures using *, ** or ***, respectively, while comparisons with a p-value > 0.05 were not marked (to reduce clutter).

### Statistical tests for assessing the significance of MVRG coefficients within one experimental condition

To determine whether there was a significant relationship between an SM tracklet property and an associated actin speckle property, a one-sample *t*-test was performed, with the null hypothesis that the single-cell MVRG coefficients followed a normal distribution with mean zero (MATLAB function ttest()). A relationship was considered significant if the t-test yielded a p ≤ 0.05. To assist with visualizing these relationships, the MVRG coefficient plots were annotated with overhead triangles, where the triangle slopes represented the significance and sign of the MVRG coefficients. Not-significant MVRG coefficients were represented with rectangles.

As additional evidence for the significance of any identified SM tracklet-speckle property relationship (where the MVRG coefficient t-test yielded a p-value ≤ 0.05), we performed MVRG analysis after dataset shuffling and calculated the fraction of shuffled data MVRG coefficients with a p-value smaller than that of the real data MVRG coefficient (**Table S7**). Specifically, suppose we were performing MVRG using *n* SM properties and *m* speckle properties. To obtain the corresponding shuffled data MVRG coefficients, the pairing of the *n* SM tracklet properties and the *m* speckle properties was shuffled, while keeping the *n* SM properties together and the *m* speckle properties together. For each MVRG analysis, the shuffling analysis was performed 100 times. The fraction of p-values from shuffled data smaller than significant p-values from real data was 0 to 0.06, further supporting the significance of the identified relationships.

To compare the MVRG coefficients between the “Free” and “Confined” subsets of an SM tracklet-actin speckle property pair for an experimental condition, a paired-sample t-test was performed (MATLAB function ttest()). A paired-sample t-test was appropriate in this case as each movie provided measurements for the “Free” subset and the “Confined” subset.

### Statistical tests for comparing MVRG coefficients between multiple experimental conditions

To assess whether MVRG coefficients (for an SM tracklet-speckle property pair) were significantly different between two experimental conditions, a two-sample one-tailed *t*-test (either right- or left-tailed, based on the observed difference), for equal means without assuming equal variances, was performed (MATLAB function ttest2()). If more than two experimental conditions were compared, multiway analysis of variance (ANOVA) with default linear model was performed (MATLAB function anovan()), followed by Tukey-Kramer post hoc analysis (MATLAB function multcompare()). Differences with a p-value ≤ 0.05 were considered significant.

## Results and Discussion

### SMI-FSM of PM proteins and CA in live single cells

To perform SMI-FSM in live single cells, we generated a stable TIME cell line (Telomerase-Immortalized Microvascular Endothelial cell line) expressing mNeonGreen-actin from a truncated CMV promoter (TIME-mNGrActin cell line). The combination of a truncated CMV promoter and cell selection through FACS ensured low labelling of actin for FSM (Fig. 1A, Video S1). We then transiently expressed in TIME-mNGrActin cells Halo-tagged versions of the surface receptor CD36 (Halo-CD36), of a chimeric transmembrane protein that binds actin (Halo-TM-ABD; positive control) or of a mutated form of TM-ABD with the mutation R579A to inhibit binding to actin (TM-ABD*; negative control) (*51*) (Fig. S2A). Labeling of the Halo-tagged proteins with JF549-conjugated Halo ligand thus enabled SMI alongside FSM (Fig. 1A, Video S2). We used for these experiments TIME-mNGrActin cells of passage 16 or higher, because at these passages TIME cells exhibited very little to no endogenous CD36 (Fig. S2B; as is the case with higher-passage primary microvascular endothelial cells (*5*)), allowing us to work exclusively with the exogenous Halo-CD36. Halo-CD36 was able to trigger downstream signaling in response to stimulation by its ligand thrombospondin-1 (TSP-1) as expected, indicating that it was functional (Fig. S2C) (*5, 52*).

**Figure 1.**
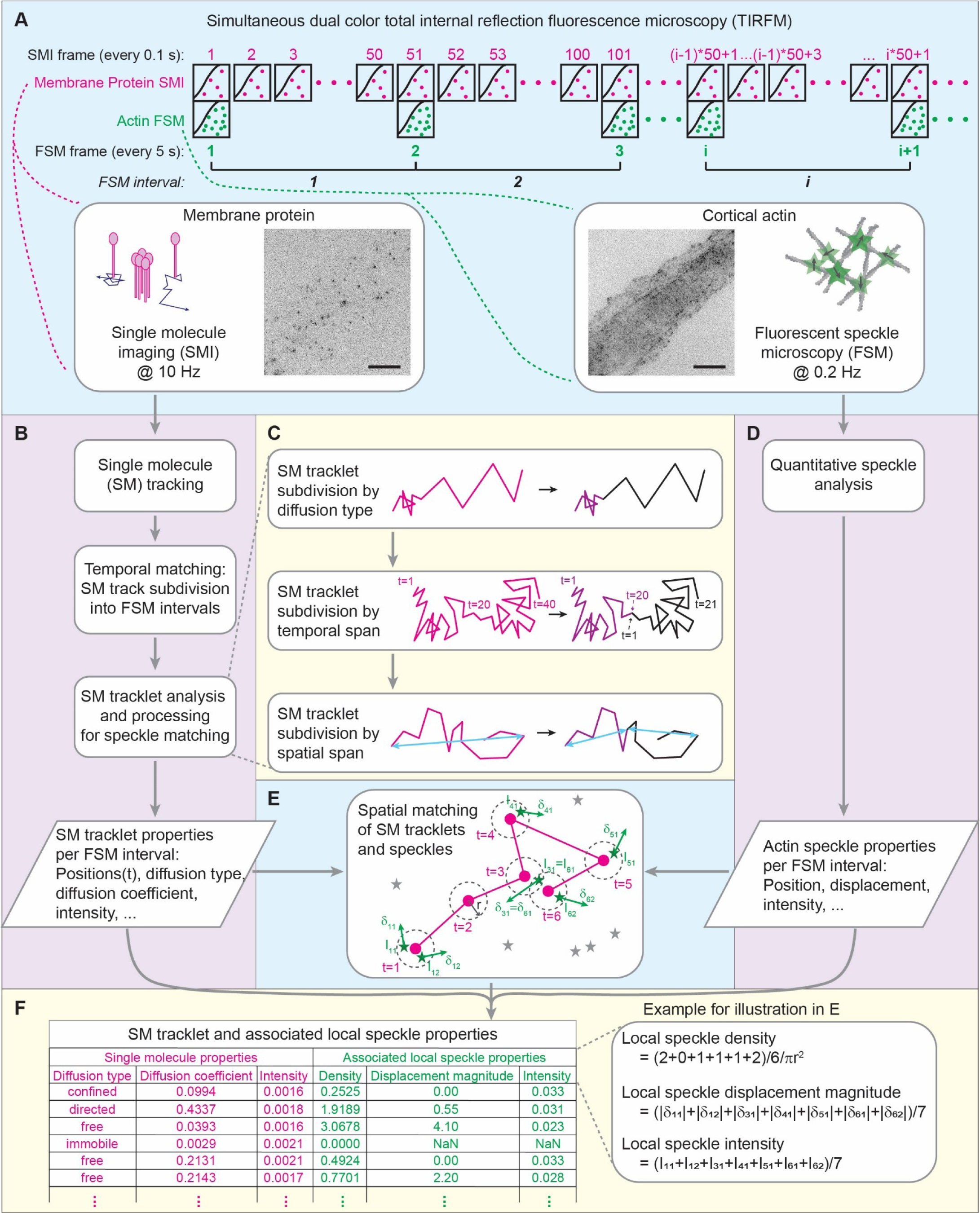
SMI-FSM yields dynamic properties of PM proteins and neighboring actin speckles. **(A)** SMI-FSM data acquisition schematic, showing the relationship between SMI and FSM frames. 51 SMI frames from FSM frame *i* to FSM frame *i*+1 correspond to FSM interval *i*. Insets, example PM protein SMI (left) and cortical actin FSM (right), with illustrations. Scale bar, 10 µm. **(B)** Major data processing steps of the SMI branch. **(C)** Breakdown of the SM tracklet analysis and processing steps. Magenta indicates a tracklet before processing; purple and black indicate tracklets after processing. t = tracklet timepoints; cyan double-arrows represent the spatial span of each tracklet. **(D)** Data processing in the FSM branch. **(E)** Illustration of spatial matching of SM tracklet (magenta dots and connecting lines) and neighboring speckles (green stars). Dashed circles = area for matching, with radius *r*. *Itm* = intensity and *δtm* = displacement (vector) of speckle *m* matched to SM at tracklet timepoint *t*. Gray stars = speckles not matched to this SM tracklet. **(F)** Example table with SM tracklet and associated local speckle properties. Inset to right: Calculation of local speckle properties for example in E.

To maximize the information gained from SMI-FSM experiments, it was important to monitor PM proteins and CA each at its relevant timescale. Thus, we performed SMI at a frame rate of 10 Hz (*7, 45*) and FSM at a framerate of 0.2 Hz (*47*) in the same cell at the same time (Fig. 1A). We used total internal reflection fluorescence (TIRF) microscopy for both, in order to focus on the bottom cell surface and the subjacent cortex. SMI videos were then analyzed using particle tracking (*42*) to obtain SM tracks of the imaged PM proteins (Video S3). At the same time, the associated FSM videos were analyzed using qFSM software (*46*) to detect and track the CA speckles (Video S4), from which various actin speckle properties were derived (see Materials and Methods for all details).

### Integration of PM protein and neighboring actin speckle properties from SMI-FSM data

In order to derive quantitative relationships between the properties of PM proteins and CA from SMI-FSM, we developed a computational pipeline linking SM tracks to their neighboring speckles in space and time, allowing us to construct a multidimensional dataset of PM protein properties and associated CA speckle properties (Fig. 1). The pipeline consisted of the following steps:

#### Step 1. Temporal matching of SM tracks and speckles

With our imaging strategy, each FSM frame-to-frame interval spanned 51 SMI frames (FSM frame 1 = SMI frame 1, FSM frame 2 = SMI frame 51, etc.; Fig. 1A). For temporal matching between SM tracks and speckles, we subdivided the SM tracks by the FSM intervals (Fig. 1B). An average of 44% of SM tracks (range: 37-49% per movie) crossed FSM interval boundaries, and were thus segmented into multiple tracklets. Henceforth, we will refer to the temporally matched SM trajectories as SM tracklets.

#### Step 2. SM tracklet analysis and processing for spatial matching with speckles

In preparation for matching (the temporally-matched) SM tracklets to actin speckles in space, we performed the following SM tracklet analyses and processing (Fig. 1C). Our goal was to obtain a collection of SM tracklets such that each tracklet (i) was as localized as possible in space to increase the accuracy of matching SM tracklets and actin speckles, and (ii) exhibited homogeneous diffusion properties to increase the sensitivity of our subsequent analysis of the coupling between SM behavior and neighboring CA speckle behavior. Note that SM track processing prior to this integration step already segmented tracks at merging and splitting events to avoid changes in clustering properties within each tracklet (described in Materials and Methods).

Step 2a. We obtained the diffusion properties of SM tracklets using transient diffusion analysis (DC-MSS (*53*)). Transient diffusion analysis allowed us to segment an SM tracklet into multiple tracklets if a change in diffusion type was detected. An average of 1.7% of SM tracklets (range: 0.7-3.6% per movie) exhibited diffusion type switches. With this, each SM tracklet (after segmentation, if any) was characterized by its diffusion type (free, confined, immobile, directed) and diffusion coefficient.

Step 2b. After Step 2a, SM tracklets could have a duration between 10 timepoints (minimum duration retained; see Materials and Methods) and 51 timepoints (number of SMI frames within an FSM interval). To reduce the variability in tracklet duration, and thus the variability in the amount of information contained in each SM datapoint when analyzing the coupling between SM behavior and actin speckle behavior, we segmented tracklets lasting ≥ 40 timepoints (twice the minimum duration possible for diffusion analysis (*53*)) in half. Each half became a separate tracklet, and its diffusion properties (diffusion type and diffusion coefficient) were calculated. An average of 3.9% of SM tracklets (range: 1.9-7.0% per movie) from Step 2a had a duration ≥ 40 timepoints.

Step 2c. Some SM tracklets, particularly those of fast-moving molecules, spanned a distance > 3 µm. At such length scales, CA properties could vary spatially (*54, 55*), and it would be meaningless to combine actin speckle properties along the tracklet for subsequent analysis steps. Thus, to obtain spatially-localized tracklets, we segmented SM tracklets spanning a distance > 3 µm into shorter daughter tracklets, each spanning a distance < 3 µm. The spatial span of an SM tracklet was defined as the maximum distance between any pair of positions along the tracklet. Any tracklet with a span between (*m*-1)*3 µm and *m*\*3 µm (where *m* = 2, 3, …) was segmented into *m* daughter tracklets of roughly equal duration. Any resulting daughter tracklets with a spatial span > 3 µm were subsequently segmented further, using the same procedure, until all tracklets in the final set had a spatial span ≤ 3 µm. An average of 3.7% of the tracklets (range: 1.0-5.6% per movie) from Step 2b spanned a distance > 3 μm. Because all SM tracklets after Step 2b had a duration < 40 timepoints, daughter tracklets resulting from this step had a duration < 20 timepoints and their diffusion properties could not be analyzed separately. Rather, the daughter tracklets from this step inherited the diffusion properties (diffusion type and diffusion coefficient) of the original SM tracklet.

With these steps, we obtained the following properties for each SM tracklet:

- Diffusion type
- Diffusion coefficient
- SM spatial span
- SM intensity, defined as the mean particle intensity over the tracklet duration.

#### Step 3. Spatial matching of SM tracklets and actin speckles

For each SM tracklet in an FSM interval, we determined its neighboring speckles based on its per-timepoint positions and the speckle positions at the starting frame of the FSM interval (Fig. 1D, E). Specifically, speckles that were within a distance *r* of 3.5 pixels (311.5 nm) from any position within the SM tracklet were considered neighboring speckles. The radius r = 3.5 pixels primarily accounted for the registration shift between the two cameras (for the two channels) in our imaging setup (*45*).

#### Step 4. Calculation of local speckle properties around SM tracklets

After matching speckles to SM tracklets, we calculated the local speckle properties around each SM tracklet. Consider an SM tracklet made of *N* timepoints. Let *s_t_* be the set of speckles, and *M_t_* be the number of speckles, matched to the SM tracklet at timepoint *t*, *t* = 1…*N*. Note that a speckle could be matched to an SM tracklet at multiple timepoints (see example in Fig. 1E), and it would contribute a measurement at each timepoint where it was matched to the SM tracklet. Defining 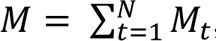, we calculated the local speckle properties around an SM tracklet as follows:

- Local speckle density *ρ*, defined as:

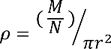

where *r* is the radius (= 3.5 pixels) for matching speckles to the SM tracklet. Co-imaging of CA speckles and phalloidin-stained actin demonstrated that local speckle density reflected local CA density (Fig. S3, **Supplemental Note 1**).
- Local speckle displacement magnitude Δ: Let 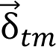 be the displacement of speckle *m* matched to the SM tracklet at timepoint *t* (*m* = 1…*M_t_*, *t* = 1…*N*). Then the local speckle displacement magnitude Δ was defined as:

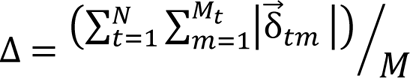 Note that local displacements with 0 magnitude were considered “not measurable” (Materials and Methods) and did not contribute displacement information for any subsequent analyses. Local speckle displacement represented network remodeling resulting from filament polymerization, depolymerization and motor protein induced movement (*48, 56*).
- Local speckle intensity *I*: Let *I_tm_* be the intensity of speckle *m* matched to the SM tracklet at timepoint *t*. Then the local speckle intensity *I* was defined as:

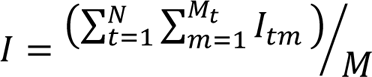 Local speckle intensity represented the quantity of fluorescently labelled actin.

Through Steps 1-4, we put together, for each SM tracklet, a list of its properties (Step 2) and its local actin speckle properties (Step 4) (Fig. 1F). SM tracklets without any matched speckles (∼20% of SM tracklets) had a zero speckle density and no other associated speckle properties. We then compiled together the SM tracklet and associated local speckle properties from all FSM intervals of an individual cell. Outliers within each property were identified and removed (using the interquartile range method, also called Tukey’s fences method (*50*), with conservative thresholds that retained about 99% of the datapoints). Altogether, this provided us with a multidimensional dataset of SM tracklet and associated local speckle properties to investigate their relationships in an explicit and quantitative manner.

### The spatiotemporal organization of PM proteins reflects their actin-binding ability

Application of the analysis pipeline to SMI data of CD36, TM-ABD and TM-ABD* revealed that ∼60% of their tracklets exhibited free diffusion, ∼20% exhibited confined diffusion, and the remaining ∼20% were immobile or exhibited directed diffusion (Fig. 2A). The actin-binding control, TM-ABD, had a significantly lower fraction of free SM tracklets, and correspondingly higher fractions of confined and immobile SM tracklets, than the actin-non-binding control, TM-ABD*. CD36 exhibited intermediate behavior between the actin-binding and non-binding controls. The diffusion coefficients per diffusion type (free or confined) of the three PM proteins were similar (Fig. 2B). Our results for TM-ABD and TM-ABD* are consistent with previous studies, where it was observed (although using different labeling and imaging techniques and on different time scales) that TM-ABD undergoes more tethering and/or slower diffusion than TM-ABD* (*15, 51*). The intermediate behavior of CD36 suggests that its actin-binding ability is weaker and less direct than that of TM-ABD, but stronger and more specific than that of TM-ABD*. Indeed, the link between CD36 and actin is most likely indirect, through intermediate molecules (*13, 57, 58*).

**Figure 2.**
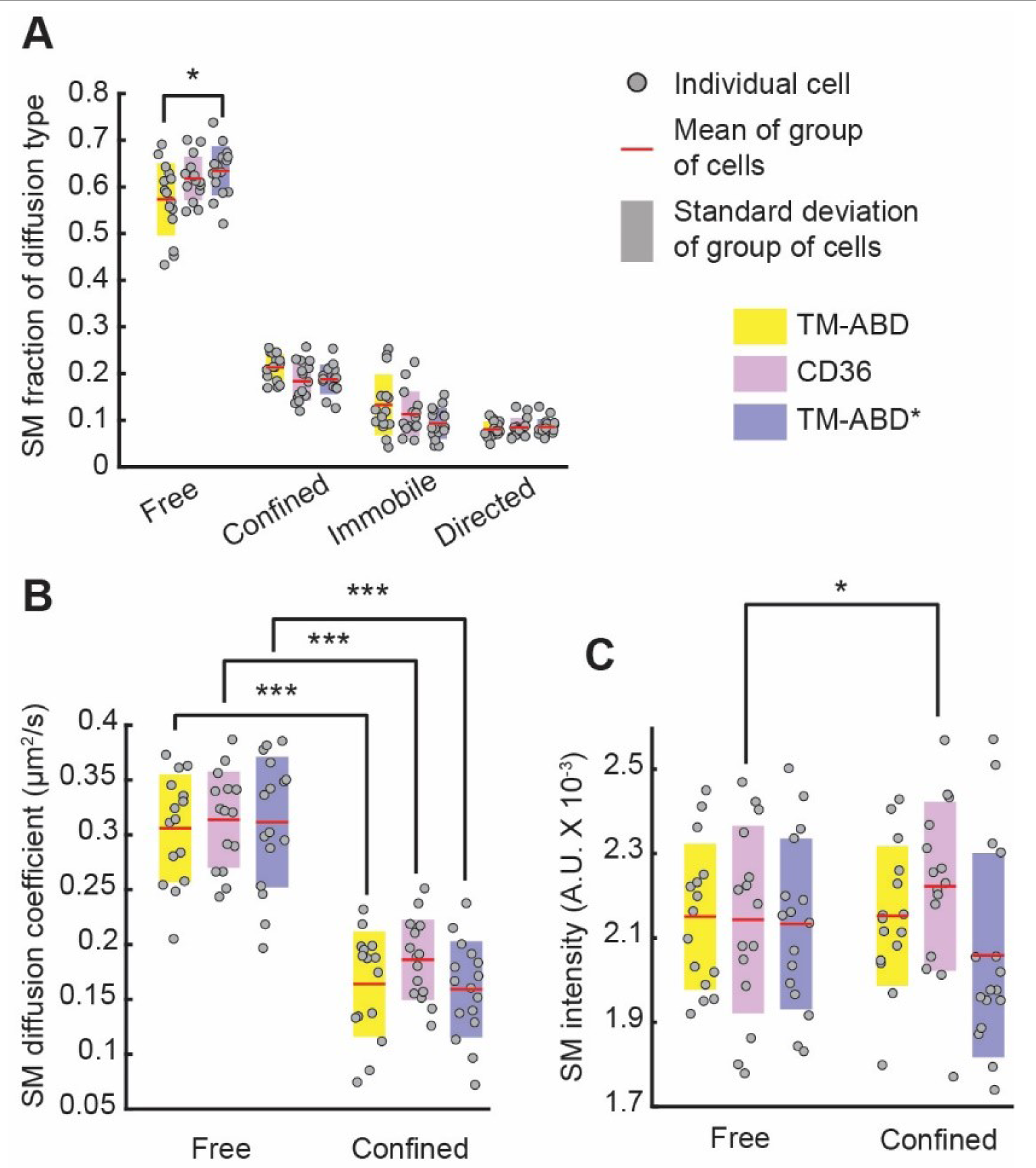
Dynamic organization of PM proteins reflects their actin binding abilities. **(A)** Fraction, **(B)** diffusion coefficient and **(C)** intensities of SM tracklets of CD36 (magenta), TM-ABD (yellow) and TM-ABD* (blue) undergoing the indicated diffusion types. Gray circles show individual cell measurements (fraction in A and average in B and C). Red lines and shaded bars show mean and standard deviation, respectively, over group of cells representing a particular PM protein. Different PM proteins were compared using ANOVA followed by Tukey-Kramer post-hoc analysis; free and confined properties for the same PM protein were compared using paired t-tests. *, ***: p-value ≤ 0.05, 0.001, respectively. p-values > 0.05 are not explicitly indicated. See Table S8 for sample size and other dataset details.

Our analysis also shed light on the clustering of the imaged molecules (captured by their SM intensity) and its relation to their diffusion type. Here (and for most subsequent analyses) we focused on the free and confined SM tracklets, as they constituted the majority of tracklets. Our analysis revealed that confined CD36 tracklets had significantly higher intensities than free CD36 tracklets (Fig. 2C). TM-ABD and TM-ABD*, on the other hand, showed no difference in SM intensity between free and confined tracklets (Fig. 2C). The unique trend exhibited by CD36 (vis-à-vis TM-ABD and TM-ABD*) reflects the functional significance of CD36 clustering, in that clustering enhances CD36-downstream adaptor interactions which could lead to confinement (*5, 52*). It is worth noting that CD36 clustering has been demonstrated to depend on actin (*5*).

### PM proteins of different diffusion types are associated with different local CA dynamics

Next, using the actin speckles that were matched to SM tracklets of CD36, TM-ABD and TM-ABD* (Fig. 1E, Fig. 3A), we investigated the local speckle properties around the different PM proteins. We reasoned that the local CA environment would have the largest influence on a PM protein’s spatiotemporal organization. While local speckle density (Fig. 3B) and intensity (Fig. 3C) were not different between different PM proteins and diffusion types, local speckle displacement magnitude was significantly higher for confined SM tracklets than for free tracklets, for all imaged PM proteins (Fig. 3D). Moreover, the difference was largest and most significant for TM-ABD, followed by CD36, and lastly TM-ABD*. Note that the corresponding cell wide (“global”) speckle properties did not differ between the cells expressing the different proteins, as expected (Fig. S4). As speckle displacement represents network remodeling (*48, 56*), these observations suggest that free/confined PM proteins are associated with regions of lower/higher CA network remodeling. Moreover, the difference in local CA environment between free and confined PM proteins is coupled to the extent of the PM protein’s binding to actin (TM-ABD > CD36 > TM-ABD*). All in all, these results highlight the high sensitivity of SMI-FSM, enabling the unique characterization of the local CA environment around PM proteins with different actin-binding ability and mobility.

**Figure 3.**
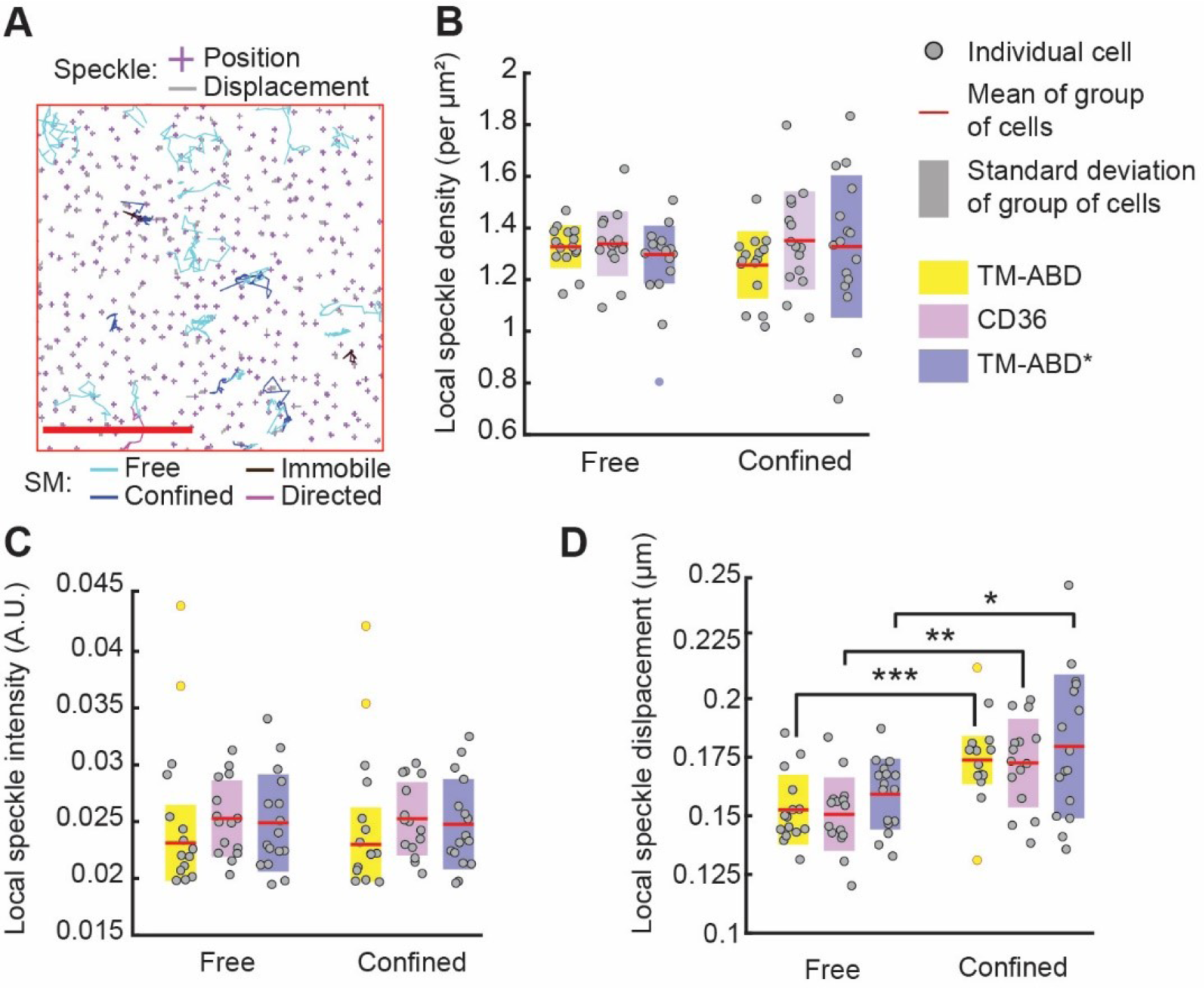
Local CA environment varies by PM protein diffusion type. **(A)** CD36 tracklets in one FSM interval overlayed on actin speckle positions at the beginning of that interval. Scale bar, 5 µm. **(B)** Density,**(C)** intensity and **(D)** displacement magnitude of speckles localized around SM tracklets of CD36 (magenta), TM-ABD (yellow) and TM-ABD* (blue) undergoing indicated diffusion types. Colors, circles, red lines, shaded bars and statistical tests as in Fig. 2. ** in D: p-value ≤ 0.01. Non-gray circles in C and D indicate outliers. See Table S8 for sample size and other dataset details.

### PM protein mobility-CA relationships vary by CA density and PM protein diffusion type

To more closely inspect the relationship between PM protein behavior and the local CA environment at the SM level, we sought to quantify the (mathematical) dependence of SM tracklet properties (e.g. diffusion coefficient or intensity) on speckle properties through linear multivariate regression (MVRG). MVRG would allow us to model the *collective* relationship between multiple CA speckle properties and SM diffusion coefficient or intensity. We utilized linear MVRG as the simplest model to quantify the coupling and characterize the relationship between these SM tracklet properties and speckle properties.

As a first step, and particularly to verify the validity of a linear relationship as modeled by MVRG, we made pairwise scatterplots of SM diffusion coefficient or SM intensity vs. various actin speckle properties (separately for free and confined SM tracklets) on an individual cell basis (Fig. S5). While most pairwise relationships appeared linear, the scatterplot of SM diffusion coefficient vs. local actin speckle density exhibited a biphasic “inverted v-shape” relationship in most cells (70% for CD36 and TM-ABD and 63% for TM-ABD* through objective assessment; **Supplemental Note 2**; Fig. S6A, B). Automatic estimation of the threshold at which the relationship switched from positive to negative (**Supplemental Note 2**) showed that it was similar between all cells and conditions, and was very close to the overall (global) speckle density in the imaged cells (1.18 vs. 1.25 speckles/µm^2^; Fig. S6C and S4A). The “inverted v-shape” relationship between SM diffusion coefficient and local speckle density may be reflective of differential influence of CA on PM protein mobility at different CA density levels (*4, 25*). However, this shape may also be a consequence of the discrete nature of speckles and spatial sampling by SM tracklets of different diffusion coefficients (**Supplemental Note 3**). In either case, to satisfy the assumption of linearity in MVRG, we distinguished between SM tracklets based on their local speckle density level: low density/sparse (local speckle density < global density) and high density/dense (local speckle density ≥ global density). This yielded four subgroups of SM tracklets for MVRG analysis, based on the combination of diffusion type (free vs. confined) and local speckle density level (sparse vs. dense) (Fig. S6D).

MVRG of SM tracklet diffusion coefficient on various CA speckle properties (local density, displacement magnitude and intensity) yielded significant MVRG coefficients for only speckle density and displacement magnitude (Fig. 4A; Fig. S7). The two speckle properties exhibited very little multicollinearity (variance inflation factor (VIF) ≤ 1.96 for 97.8% of analyzed cells), allowing us to use them as independent predictors of SM tracklet diffusion coefficient (*49*). The MVRG coefficients with respect to local speckle density were positive for the low density subgroups and negative for the high density subgroups, as expected from the above biphasic analysis. The MVRG coefficients with respect to local speckle displacement magnitude were universally negative (or, in the case of TM-ABD* at low density, not significant), indicating a negative relationship between PM protein diffusion and CA remodeling. This relationship was consistent with our earlier observation that higher mobility diffusion types are associated with regions of lower CA network remodeling (Fig. 3D).

**Figure 4.**
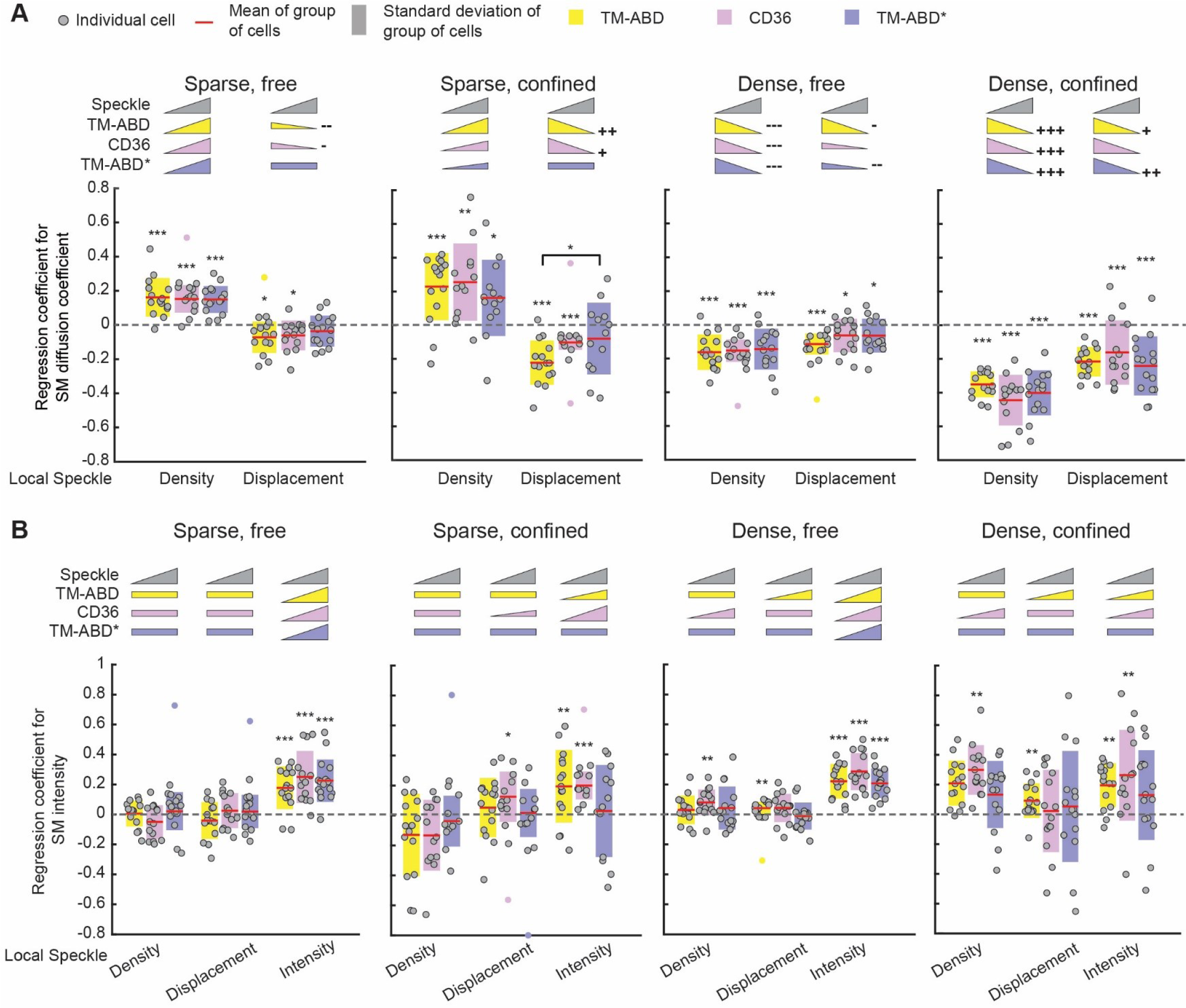
MVRG analysis reveals multiple regimes of PM protein-CA relationships. Coefficients from MVRG of **(A)** SM diffusion coefficient on local speckle density and displacement magnitude, and **(B)** SM intensity on local speckle density, displacement magnitude and intensity for CD36 (magenta), TM-ABD (yellow) and TM-ABD* (blue). MVRG analysis was performed separately for the four indicated subgroups of SM tracklets based on diffusion type and local speckle density pairings. Colors, circles, red lines and shaded bars as in Figs. 2 and 3. MVRG coefficients significantly different from zero (dashed grey line), tested by t-test, are indicated by *, **, *** above the MVRG coefficient plots for p-value ≤ 0.05, 0.01, 0.001, respectively. Differences between the MVRG coefficients of different PM proteins, tested by ANOVA followed by Tukey-Kramer post-hoc analysis, are indicated by * above bracket for p-value ≤ 0.05. Inset pictograms: visual summary of MVRG results. Triangle slope and direction represent significance and sign (positive or negative) of MVRG coefficient; rectangles indicate not significant MVRG coefficients. Next to the triangles, +, ++, +++ (and corresponding -, --, ---) indicate significant differences between free and confined MVRG coefficients at the same density level (e.g., sparse free vs. sparse confined) for each PM protein, tested by paired t-test, for p-value ≤ 0.05, 0.01, 0.001, respectively. In all tests, p-values > 0.05 are not explicitly indicated. See Table S8 for sample size and other dataset details.

All in all, MVRG analysis yielded the following insight into the relationship of CA properties to PM protein diffusion coefficient: (i) Integrating across the four subgroups, the actin binding control, TM-ABD, showed the strongest relationships with CA (i.e., most significant regression coefficients), the actin non-binding control, TM-ABD*, showed the weakest relationships, and CD36 was in between. In other words, the strength of the identified relationships reflected the actin binding ability/strength of the three PM proteins. (ii) At either density level, confined tracklets showed overall stronger relationships with CA than free tracklets, especially for TM-ABD and CD36, suggesting that confined PM proteins are more strongly associated with CA than freely diffusing PM proteins. (iii) Overall, free tracklets associated with low speckle density had the weakest relationships with CA, confined tracklets associated with high density had the strongest, while the other two subgroups were in between. (iv) Interestingly, differences in CA-PM protein relationships between the three proteins were detected for the diffusion type-speckle density subgroups with weaker relationships. This suggests that regimes of weaker CA influence allow for differential regulation of PM protein mobility in a manner that reflects their actin binding ability/strength, in contrast to regimes of strong and ubiquitous CA influence (as is the case of confined tracklets associated with high speckle density, where all PM proteins exhibit the same relationship with CA). These results highlight the ability of SMI-FSM to reveal the quantitative relationships between CA and PM protein dynamics, and its sensitivity to identify differences between them reflecting their actin binding ability.

### PM protein clustering exhibits relationships with CA only at high CA density

We next performed MVRG analysis for SM tracklet intensity (proxy for PM protein clustering). For consistency with the above analysis, we performed the analysis separately for the four subgroups of diffusion type-speckle density combinations. In the case of SM tracklet intensity, local speckle density, displacement and intensity exhibited significant MVRG coefficients (Fig. 4B). As above, the three speckle properties exhibited negligible multicollinearity with each other (VIF ≤ 2.5 for density, ≤ 3.2 for displacement, and ≤ 3.01 for intensity in 97.8% of analyzed cells) (*49*). The MVRG coefficients with respect to intensity were significant and positive under all conditions, except for confined TM-ABD* tracklets. This uniformity in the MVRG coefficients with respect to speckle intensity, in addition to the speckle intensity’s negligible multicollinearity with other speckle properties, suggests that the positive relationship between SM and speckle intensities reflects systematic variations in intensity in both channels, due to slight illumination heterogeneities and/or undulations in the position of the bottom cell surface. Including speckle intensity in the MVRG analysis accounted for this “artifactual” source of intensity variation in the SM tracklets, allowing us to then uncover the relationship between SM tracklet intensity and other local speckle properties.

The MVRG coefficients with respect to the other relevant local speckle properties, namely density and displacement, varied by PM protein type and actin density level. At high density, CD36 clustering showed a significant positive relationship with actin density, consistent with our previous observation that actin rich regions support higher-order CD36 clustering (*5*). TM-ABD clustering, on the other hand, showed a significant positive relationship with local speckle displacement at the high CA density level. This is consistent with previous observations that clustering of the ABD of ezrin requires actin movement, with no dependence on actin concentration (*59*). These relationships between CD36 or TM-ABD tracklet intensity and speckle properties were observed only at the high density level; the MVRG coefficients at the low density level were largely not significant. Not surprisingly, TM-ABD* did not show any relationship with either speckle density or displacement at any time, as expected given its inability to bind actin (*25, 59*).

Altogether, these results further demonstrate the ability of SMI-FSM to accurately identify the relationships between the dynamic organization of PM proteins and CA, and sensitivity to distinguish between PM proteins with different mechanisms of actin-mediated movement and clustering. As with PM protein mobility, PM protein clustering also exhibited differential relationships (or lack thereof) with CA based on CA density level. This highlights the need to study the relationships between PM proteins and CA in cellular settings that preserve dynamic information and spatial context.

### Response of PM proteins to actin perturbation varies based on local actin environment

Cell wide actin perturbation is a commonly used approach to investigate the role of actin in regulating PM protein spatiotemporal organization. An inherent assumption in such approaches is that actin perturbation, and the corresponding PM protein modulation, are homogeneous throughout the cell. Given our observations so far that the local CA environment, and its relationship to PM protein mobility and clustering, vary by CA density level and PM protein diffusion type, we sought to test the validity of this assumption. SMI-FSM is ideally suited for this task, as it allows the study of global CA architecture and dynamics as well as their local counterparts near individual PM proteins, and the study of PM proteins throughout the whole cell and at different CA density levels.

For these experiments, we used Halo-tagged CD36 as our model PM protein. For actin perturbation, we used a mild concentration (30 nM) of Lat A, in order to maintain overall cell morphology for TIRF imaging (Videos S6, S7). Lat A depolymerizes actin and causes severing of actin filaments (*60*). Here, to monitor the effects of Lat A over time, we devised a data acquisition scheme where we added Lat A to cells at the microscope, and then imaged the same cell at three instances (each for 1.5 min): (i) right after Lat A addition (0-2 min), 7-10 min after Lat A addition, and 14-16 min after Lat A addition (Fig. 5A). This scheme allowed us to monitor the same cell over timescales relevant for Lat A perturbation while minimizing photobleaching. As expected, Lat A decreased global CA speckle density and (gradually) increased global actin speckle displacement (Fig. 5B, C).

**Figure 5.**
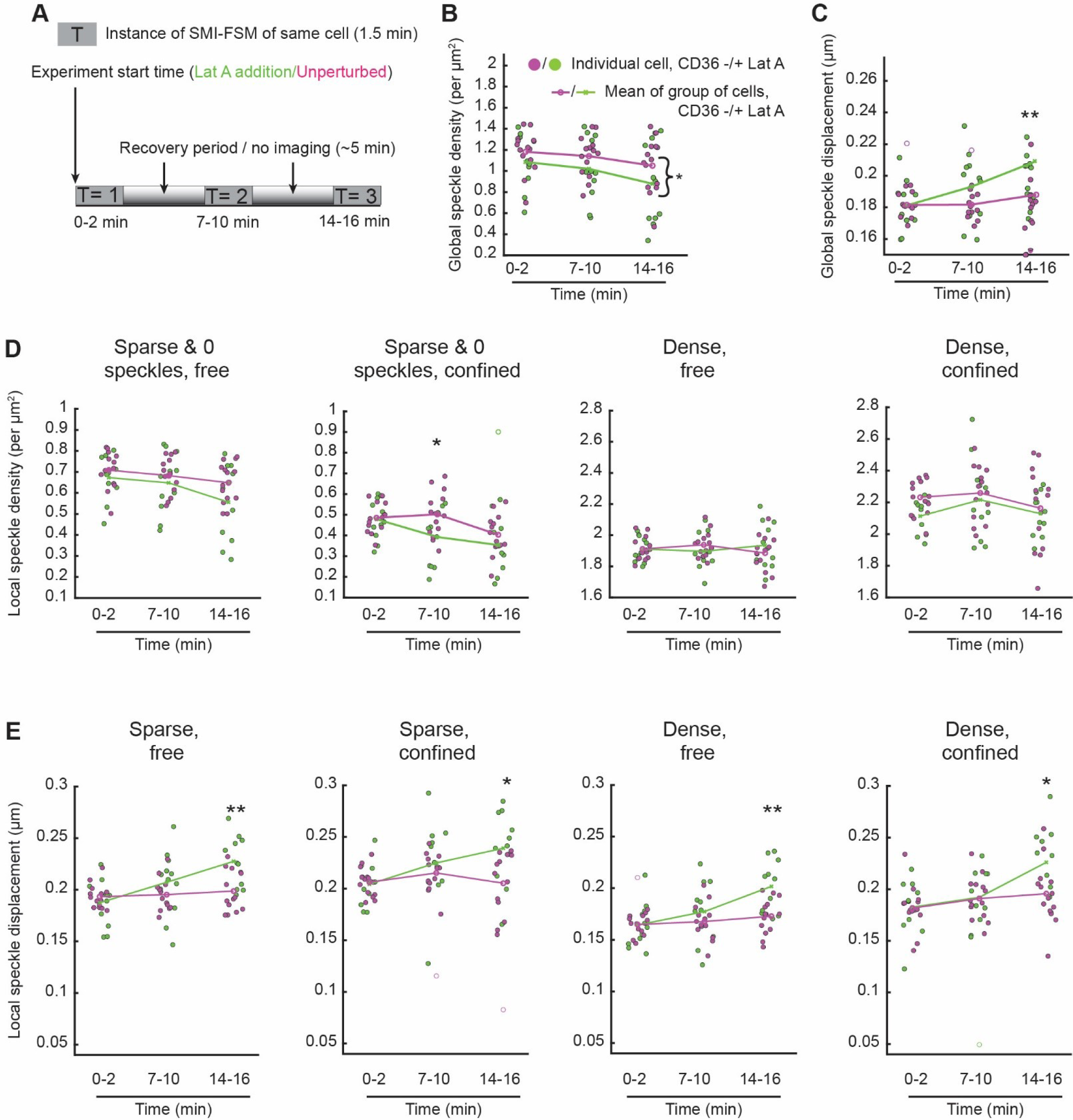
Latrunculin A treatment leads to different changes in actin speckle properties globally vs. locally around individual CD36 SM tracklets. **(A)** Schematic of SMI-FSM at multiple instances of a single cell, with recovery periods in between, in the presence or absence of Lat A. **(B, C)** Global speckle density **(B)** and global speckle displacement magnitude **(C)** of all actin speckles within the cell mask in the absence (magenta) or presence (green) of Lat A, at indicated instances. Filled circles, individual cell values from each instance. Unfilled circles with same outline as filled circles indicate outliers. The means over the inlier cells belonging to each instance are connected with lines for visual aid. Measurements from CD36 -LatA and CD36 +LatA per instance were compared by 2-sample t-test. Measurements from CD36 -LatA and CD36 +LatA across all instances were compared using a paired t-test (curly brackets). *, **: p ≤ 0.05, p ≤ 0.01. Comparisons with a p-value > 0.05 are not explicitly indicated. **(D-E)** Local speckle density **(D)** and local speckle displacement magnitude **(E)** associated with the indicated subgroups of CD36 SM tracklets, based on diffusion type and local speckle density pairings, at indicated instances. In D, “Sparse + 0 speckles” combines SM tracklets associated with low local speckle density (sparse) and SM tracklets not matched to speckles at all (extremely sparse). Colors, circles, lines and statistical tests as in B, C. See Table S8 for sample size and other dataset details.

First, we examined changes in global vs. local speckle properties upon Lat A perturbation. We found that, unlike the global speckle density, local speckle density around confined or free CD36 SM tracklets belonging to either speckle density level did not show a consistent reduction in speckle density (Fig. 5D). Of note, for speckle density calculations, SM tracklets with 0 matched speckles were combined with those associated with low speckle density. Local speckle displacement followed the global trend, although with a delay around confined SM tracklets. The Lat A vs. unperturbed lines started separating only at the third imaging instance for confined SM tracklets, while they started to separate at the second imaging instance for free tracklets and globally; Fig. 5E vs. C). These observations demonstrate that global changes in actin architecture and dynamics upon perturbation may not reflect the local CA changes around specific PM proteins. Second, we examined changes in CD36 mobility in response to Lat A perturbation for all SM tracklets vs. those of specific diffusion types or CA density levels. When putting together all density levels (“overall”), the diffusion coefficient of confined SM tracklets was higher in Lat A-treated cells in the 0-2 min instance, but then was not different from untreated cells at later instances (Fig. 6A top). The overall diffusion coefficient of free tracklets (Fig. 6A bottom) and the fraction of tracklets belonging to the different diffusion types (Fig. S8A) were similar in the absence and presence of Lat A at all instances. The observation that only the diffusion coefficient of confined tracklets was sensitive to actin perturbation is consistent with the stronger MVRG relationships observed above for confined tracklets vs. free tracklets (Fig. 4A).

**Figure 6.**
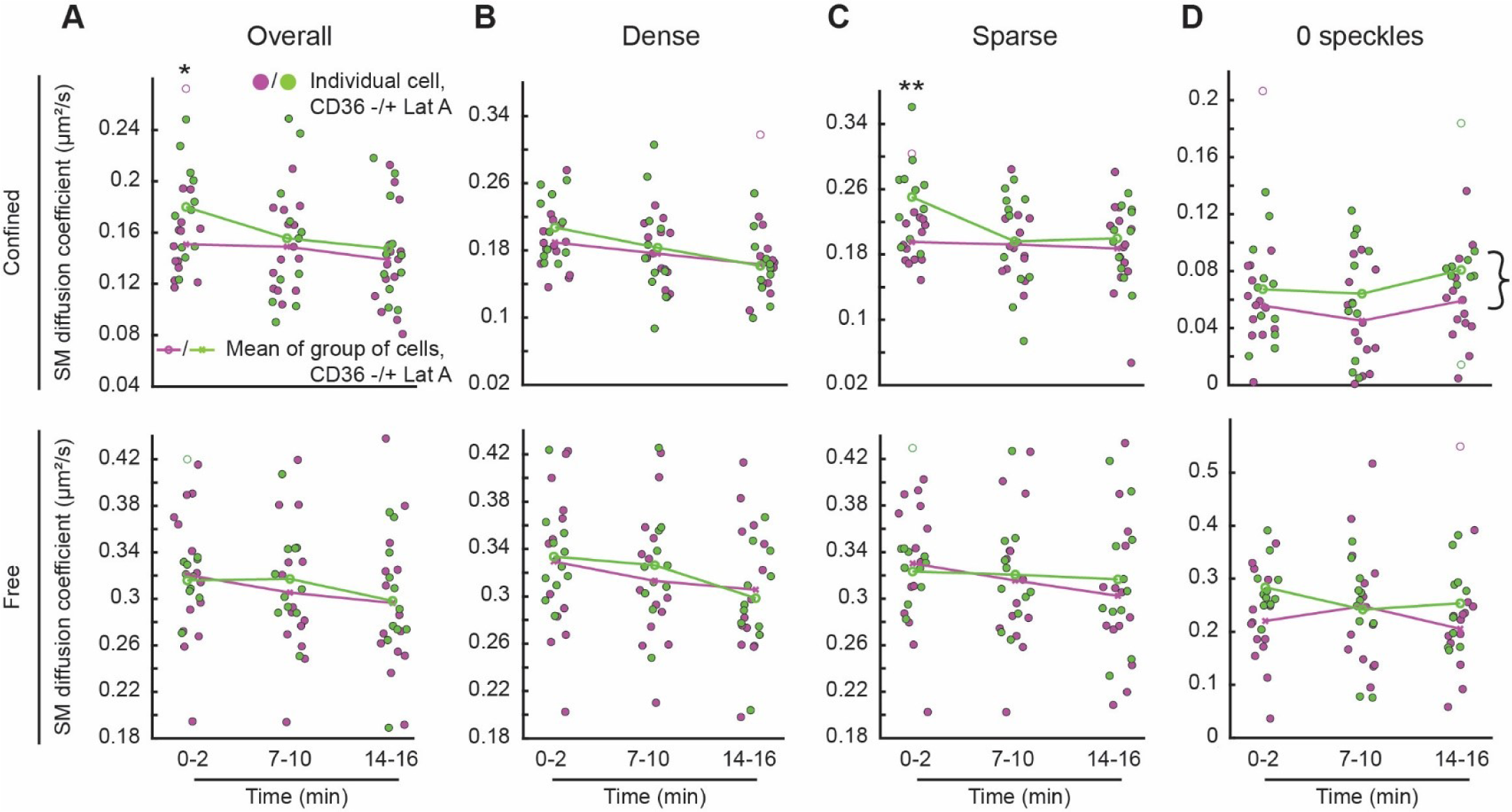
Response of CD36 mobility to Latrunculin A treatment varies by tracklet diffusion type and local actin environment. (A-D) CD36 SM diffusion coefficient in the absence (magenta) or presence (green) of Lat A, at indicated instances, as calculated for all tracklets (“overall”; **A**) and the subsets of tracklets associated with high **(B)** and low **(C)** speckle densities, and 0 speckles **(D)**. Colors, circles, lines and statistical tests as in Fig. 5. Top row, confined tracklets; bottom row, free tracklets. See Table S8 for sample size and other dataset details.

Further subdivision of the SM tracklets by their local speckle density level (Fig. S8B) revealed stark differences in the diffusion coefficient of confined tracklets (but not free tracklets) between the different density levels (Fig. 6B-D, top for confined, bottom for free). For SM tracklets associated with high local CA density, the diffusion coefficient of confined tracklets was not different between Lat A treated and untreated cells (Fig. 6B). For those associated with low CA density, the diffusion coefficient of confined tracklets was higher at 0-2 min after Lat A addition, but then similar in the absence or presence of Lat A at later instances (Fig. 6C). Interestingly, for tracklets associated with extremely low CA density (0 speckles; Fig. S3E), confined tracklets showed a significant increase in their diffusion coefficient at all instances upon Lat A treatment (Fig. 6D). As Lat A reduced global speckle density (Fig. 5B), we were concerned that the increase in diffusion coefficient for confined tracklets matched to 0 speckles could be due to the ability of SM tracklets to span a larger space without encountering any speckles. However, the span of confined tracklets matched to 0 speckles was similar in the absence and presence of Lat A (Fig. S8C), thus eliminating this concern. Thus, these observations indicate that Lat A treatment increased CD36 tracklet mobility, as expected, but in a diffusion type, CA density level and time-dependent manner.

These results highlight the complexity of PM protein responses to actin perturbations, such as the Lat A treatment performed here. It is interesting that the most sustained difference in the diffusion coefficient of confined tracklets between Lat A treated and untreated cells was at extremely low CA density (0 speckles), where the direct influence of actin on PM proteins is expected to be minimal. For these tracklets, the change in diffusion coefficient most likely reflects an overall modulation of PM organization upon actin perturbation (*37*). In contrast, for confined tracklets at the sparse and dense levels, which exhibited increasingly stronger relationships between CA and PM protein mobility (Fig. 4), the diffusion coefficient either changed transiently (sparse) or did not change at all (dense). This, together with the observation that local CA changes around individual PM proteins only partially followed the global CA changes, suggests spatial and temporal adaptation of the local CA environment and its relationship with PM proteins upon actin perturbation in order to minimize the effect of actin perturbation on PM protein behavior (at least upon mild actin perturbation). These analyses demonstrate the unique insight that SMI-FSM provides into the spatiotemporal co-organization of CA and PM proteins and its modulation upon cellular perturbations.

## Conclusions

We developed SMI-FSM, a multiscale imaging and computational analysis framework that probes the dynamic properties of both PM proteins and CA and quantifies their relationships in live single cells. SMI-FSM has enabled, for the first time to the best of our knowledge, the characterization of CA architecture and dynamics and their relationships to the spatiotemporal organization of PM proteins at high resolution. Different FSM acquisition frame rates or degrees of actin labeling (e.g., down to SM speckles (*61*)) would allow the probing of CA at different spatial and temporal scales, thus allowing the study of the multi-faceted influence of CA on PM protein organization (*21, 25*). The multidimensional dataset of PM protein and CA properties derived simultaneously from a single cell by SMI-FSM enables the exploration of the relationships between these properties, in unperturbed cells as well as upon perturbation. In our study, SMI-FSM revealed differential relationships between CA and PM proteins based on the PM proteins’ actin binding ability, diffusion type and local CA density. The analysis enabled by SMI-FSM also highlighted the complexity of cell wide actin perturbation, such that global changes in actin properties caused by perturbation were not reflected in the CA properties near PM proteins, and the change in PM protein properties upon perturbation varied based on the local CA environment. Given the widespread use of SMI as a method to study PM constituents, we expect SMI-FSM to be widely applicable to exploring the influence of CA architecture and dynamics on different PM proteins (and lipids). The analysis strategies within SMI-FSM can be also expanded, based on the questions of interest. SMI-FSM will be particularly powerful to study PM-CA interplay in the context of conditions involving actin-dependent dynamic cellular process, such as cell migration.

## Software availability

The Matlab code implementing the computational analysis pipeline of SMI-FSM is available on GitHub (https://github.com/kjaqaman/SMI-FSM).

## Supporting material

The supporting material of this manuscript consists of Document S1 (containing Supplemental notes 1-3, Figures S1-S9 and Tables S1-S8) and Videos S1-S7.

## Author contributions

A.D., H.-T.N. and K.J. designed research. A.D., B.R.A. and N.T. performed experiments. H.-T.N., A.D., D.T. and K.J. contributed to software development. A.D and H.-T.N. analyzed data. A.D, H.-T.N. and K.J. wrote manuscript with input from all authors.

## Acknowledgments

We thank the Danuser lab at The University of Texas Southwestern Medical Center (UTSW) for help with qFSM and the Mayor lab at National Centre for Biological Sciences (India) for providing the GFPTM-Ez-AFBD and GFPTM-Ez-AFBD* constructs. We thank Dr. Tieqiao Zhang for microscopy support. We also thank the UTSW BioHPC facility for providing high-performance computing systems. This work was supported by funding from the National Science Foundation (MCB-2114417), the National Institutes of Health/National Institute of General Medical Sciences (R35 GM119619) and the UTSW Endowed Scholars Program to KJ. DT was supported by a Green Fellowship (UTD/UTSW) and SURF program (UTSW).

## Video Legends

**Video S1.** FSM of CA visualized as mNeonGreen-Actin speckles at 0.2 Hz in a TIME-mNGrActin cell. This time lapse consists of 11 FSM frames = 50 s (= 10 FSM intervals). Image size is 38.9 × 32.4 μm^2^.

**Video S2.** SMI of Halo-JF-549-CD36 at 10 Hz in the same TIME-mNGrActin cell as in Video S1. This video consists of the 49 SMI frames within one FSM interval and excludes the first and last SMI frames of the interval (which coincide with the FSM frames). Image size is 38.9 × 32.4 μm^2^.

**Video S3.** SM particle tracking. SMI streams (at 10 Hz) of Halo-JF-549-TM-ABD (left), Halo-JF-549-CD36 (middle) and Halo-JF-549-TM-ABD* (right) with their respective tracks overlaid on the fluorescence images. The streams span 1 FSM interval, similar to Video S2. Random track colors are used to help distinguish between neighboring tracks. Image size is 8.9 × 10.7 μm^2^. The displayed areas are zoomed in portions of their respective images (45.5 × 45.5 μm^2^), for visual clarity. The areas are completely within the cell.

**Video S4.** FSM speckle detection and tracking. FSM time lapse (at 0.2 Hz) of mNeonGreen-Actin speckles with the speckle detections and tracks overlaid on the fluorescence image. Tracks are shown as red lines. Circles and triangles represent primary and secondary speckle detections (Mendoza, Michelle C et al. “Quantitative fluorescent speckle microscopy (QFSM) to measure actin dynamics.” Current protocols in cytometry vol. Chapter 2 (2012)). Image size is 8.9 × 10.7 μm^2^. The displayed area is a zoomed in portion of the acquired image (45.5 × 45.5 μm^2^), for visual clarity. The area is completely within the cell.

**Video S5.** Overlay of CD36 SM tracklets per FSM interval on actin speckle positions at the beginning of each interval. Speckle positions are shown as plus signs, and speckle frame-to-frame displacements as connecting lines on top. SM tracklets are color-coded by diffusion type: black – immobile, dark blue – confined, cyan – free, magenta – directed. Cell same as in Videos S1 and S2.

**Video S6.** FSM of CA visualized as mNeonGreen-Actin speckles at 0.2 Hz in a TIME-mNGrActin cell. The cell is treated with 30 nM LatA and imaged at three instances post-treatment: 0-2 min (left), 7-10 min (middle) and 14-16 min (right). Each time lapse consists of 19 FSM frames. Each image size is 26.34 × 27.23 μm^2^. The displayed area is a cropped portion of the acquired image (45.5 × 45.5 μm^2^), for visual clarity.

**Video S7.** FSM of CA visualized as mNeonGreen-Actin speckles at 0.2 Hz in a TIME-mNGrActin cell. The cell is unperturbed but imaged at three instances equivalent to Lat A-treated cells (Video S6). Each tme lapse consists of 19 FSM frames. Each image size is 45.5 × 45.5 μm^2^.

## Supporting documents

### Supplemental Note 1. Correlation of speckle density and phalloidin intensity

To interpret speckle density, we imaged actin speckles (using mNeonGreen-Actin) together with continuous staining of actin using phalloidin (conjugated to AF 647) in fixed TIME-mNGrActin cells. With its continuous staining, phalloidin was a faithful reporter of the quantity, or density, of actin in the cortex. Although the cells were fixed in these experiments, we acquired 5 FSM frames in order to detect and track the speckles and thus discard “ghost speckles,” i.e. speckles that exist for one frame only, as done for SMI-FSM experiments. Then, to investigate whether actin speckle density reflected CA density, we calculated the correlation between speckle density and phalloidin intensity, as follows (Fig. S3):

<u>Step 1.</u> We divided the imaged cell area into bins of size 15×15 pixel^2^ (1.2×1.2 µm^2^). This bin size was small enough to capture spatial heterogeneity (each bin covered the average span of an SM tracklet), while being large enough to yield a good range of number of speckles per bin.

<u>Step 2.</u> We calculated the mean phalloidin intensity within each bin (Fig. S3A, C).

<u>Step 3.</u> We calculated the number of speckles lying within each bin (Fig. S3B, D). For this, the speckle positions were taken from the first FSM frame of the actin FSM movie, after removal of ghost speckles.

<u>Step 4.</u> We calculated the correlation coefficient between phalloidin intensity and number of speckles per bin on an individual cell basis (Fig. S3E, F) (MATLAB function corrcoef()).

### Supplemental Note 2. Assessment of biphasic relationship between SM tracklet properties and associated local speckle properties

As seen in the scatterplot of Fig. S5A, the relationship between SM diffusion coefficient and local speckle density appeared to exhibit a biphasic “inverted v-shape” relationship, thus violating the linearity assumption in MVRG analysis. To objectively assess whether the relationship between any SM tracklet property and any local speckle property was biphasic, we performed the following fits and tests:

#### Step 1. Fit the data with one line

The 1-line fit (Fig. S6A, black line) was performed on an individual cell basis, using MVRG as described in Materials and Methods, but without the normalization step. Thus, the regression design matrix was the local speckle property of interest (e.g. density) concatenated with a vector of ones to account for the y-intercept of the fit. This step yielded the 1-line fit regression coefficients and residuals per cell.

#### Step 2. Determine the range of thresholds to test fitting the data with two lines

The 2-line fit required a threshold (in the speckle property value) that would split the datapoints into two groups that would be fit separately (with 1 line each). To determine this threshold, on an individual cell basis, we tested a range of thresholds. This range of thresholds was determined empirically for each group of cells of a particular condition, to ensure that sufficient data on each side of the tested thresholds was available for regression in every cell. This was done as follows: <u>Step 2a.</u> For each cell, the minimum and maximum thresholds to be tested were taken as the 15^th^ and 85^th^ percentiles, respectively, of the speckle property values in that cell.

<u>Step 2b.</u> The smallest minimum, and largest maximum, among the individual cells were then selected as the overall minimum, and overall maximum, respectively, of the threshold range for the group of cells.

#### Step 3. Fit the data with two lines

The 2-line fit (Fig. S6A, red lines) was performed on an individual cell basis, yielding for each cell the optimal threshold for splitting its data and the regression results for the data points on each side of its optimal threshold. This was achieved as follows, for each cell:

<u>Step 3a.</u> Going over the range of thresholds from the overall minimum to the overall maximum in steps of 0.0005 (e.g., for speckle density, this corresponds to 0.0005 speckles/pixel^2^ = 0.0631 speckles µm^2^), the data points of the cell were split into two groups by each threshold, and MVRG was performed for each group separately. As in the 1-line fit, MVRG was performed without the normalization step. This step yielded, for every tested threshold within the range, the regression coefficients and residuals for the two groups of data points (below and above the threshold).

Step 3b. To determine the optimal threshold among those tested, the residuals of the different fits (each corresponding to a different threshold) were compared. Specifically, for each threshold, the residuals of its fits to the data points below and above the threshold were concatenated and their variance was calculated as:

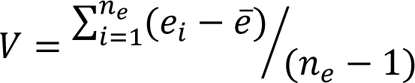

where *n_e_* is the number of datapoints, *e_i_* is the residual of datapoint *i*, and *ē* is the mean of the data points’ residuals. The optimal threshold for the 2-line fit for a cell was then taken as the threshold that minimized *V*, as the lines fit to its left and right modeled the data best. Altogether, these operations yielded the 2-line fit optimal threshold, regression coefficients, and residuals per cell.

#### Step 4. Determine whether 2-line fit or 1-line fit describes the data better

This was achieved by comparing the two fits for each cell through their Bayesian Information Criterion (BIC) values, given by:

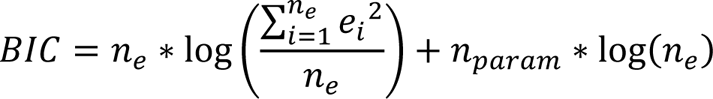

where *n_param_* is the number of parameters used in each model (and all other symbols are as in Step 3). For the 1-line fit, *n_param_* = 2. For the 2-line fit, *n_param_* = 4. For each cell, the fit with the lower BIC was taken as the better fit.

#### Step 5. Compare 2-line fit vs. 1-line fit of shuffled data to real data

As additional evidence for the validity of the 2-line fit for the real data (as assessed through Steps 1-4 above), we shuffled the SM property-speckle property matching in each cell, and repeated the 1-line and 2-line fits for the shuffled data (Fig. S6B). For the 2-line fit, the optimal threshold from the real data (for each cell) was used. This shuffling was performed 100 times for each cell, and for each shuffle we used the BIC to determine whether the 1-line fit or 2-line fit modeled the shuffled data better (Step 4).

Combining Steps 1-5, an SM property-speckle property pair were considered to exhibit a biphasic “inverted v-shape” relationship in a particular cell if they satisfied the following criteria:

1. The 2-line fit had a lower BIC than the 1-line fit.
2. The regression coefficient (line slope) was positive for the data points below the (individual cell optimal) threshold and negative for the data points above the threshold.
3. The fraction of shuffled data passing criteria 1 and 2 ≤ 5% (5 out of 100 shuffles).

With these criteria, the relationship between SM diffusion coefficient and local speckle density (see, e.g., Fig. S5A, SM diffusion coefficient vs. local speckle density) was found to be biphasic in 70% of cells expressing CD36 and TM-ABD and in 63% of cell expressing TM-ABD* (as reported in Results). No other tested SM property-speckle property pair exhibited such a systematic biphasic relationship. For example, the relationship between SM diffusion coefficient and local speckle displacement magnitude (see, e.g., Fig. S5B, SM diffusion coefficient vs. local speckle displacement) was found to be biphasic in < 7% of cells. Therefore, for MVRG analysis, the data were split into two groups based on local speckle density to satisfy the linearity assumption, but all other relationships were probed without any further data splitting.

### Supplemental Note 3. Relationship between SM diffusion coefficient, sampled area and local speckle density

The biphasic “inverted v-shape” relationship between SM diffusion coefficient and local speckle density could be the result of the combination of two factors. First, on the SM side, the area spanned and sampled by an SM tracklet was on average proportional to its diffusion coefficient (Fig. S9A). Second, on the speckle side, the discrete nature of speckles could be thought of as a “checkerboard” pattern of zeros and ones, i.e., no speckle and one speckle regions. Putting these two factors together provides an explanation for the biphasic “inverted v-shape” relationship between SM diffusion coefficient and local speckle density. Specifically, an SM tracklet with a lower diffusion coefficient, which would span a small area, would most likely get associated with only one or the other region of the “checkerboard” pattern throughout its duration. This would lead to either low or high local speckle density. On the other hand, an SM tracklet with a high diffusion coefficient, which would span a large area of the “checkerboard” pattern, would experience a combination of zeros and ones throughout its duration, and would thus get an in-between density. Indeed, when plotting SM diffusion coefficient vs. local speckle density, and color-coding the individual data points by their SM span, this trend seemed to hold (Fig. S9B).

---

**Figure S1.**
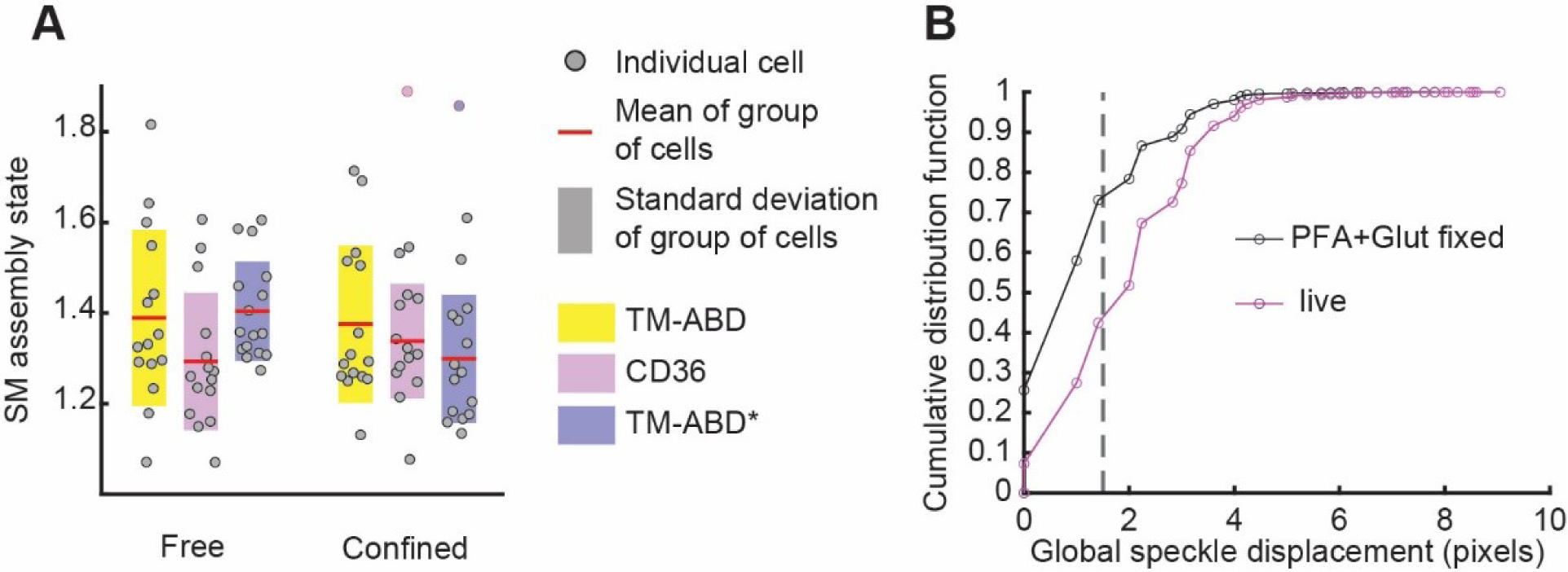
Controls for SMI conditions and FSM speckle displacement analysis. **(A)** SM assembly state. Colors, circles, red lines, shaded bars and statistical tests as in Figs. 2 and 3. **(B)** Cumulative distribution function of individual speckle displacement magnitudes derived from all speckles pooled from all cells per indicated condition. Magenta and black solid lines indicate live and glutaraldehyde-fixed cells, respectively. Dashed black line represents the threshold, 1.5 pixels, below which speckle displacements are considered artifactual (see Materials and Methods for details). See Table S8 for sample sizes and other dataset details.

**Figure S2.**
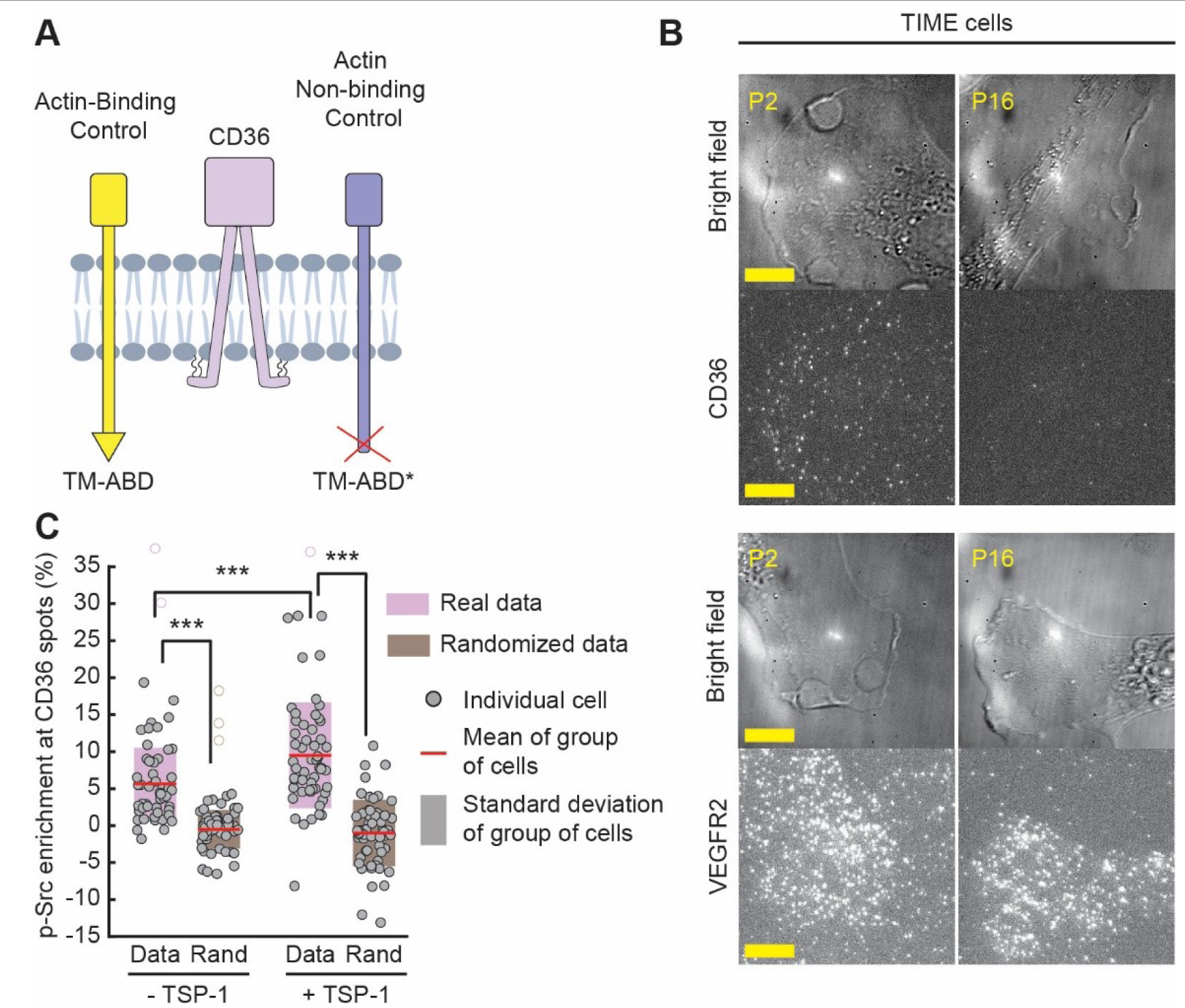
Schematic of PM proteins and characterization of CD36 expression and activity in TIME cells. **(A)** Schematic of the PM proteins investigated. **(B)** Images of CD36 and VEGFR2 endogenous expression levels at different passages of TIME cells. Bright field images are shown above each fluorescence image for context. Shown images are representatives of 5-6 cells at each passage. Scale bar: 10 µm. **(C)** phospho-Src (p-Src) enrichment at CD36 spots in the absence or presence of the CD36 ligand thrombospondin-1 (TSP-1), determined using TIRFM and signal enrichment analysis (see Materials and Methods for details). Circles, red lines and shaded areas as in Figs. 2 and 3. ANOVA followed by Tukey-Kramer post hoc analysis was used to compare between the 4 subgroups (Data/Randomized, with/without TSP-1), ***, p ≤ 0.001. Comparisons with a p-value > 0.05 are not explicitly indicated. Note that -TSP-1 Data vs. +TSP-1 Rand and +TSP-1 Data vs. -TSP-1 Rand exhibit significant differences, but they are not shown to reduce clutter. See Table S8 for sample size and other dataset details.

**Figure S3.**
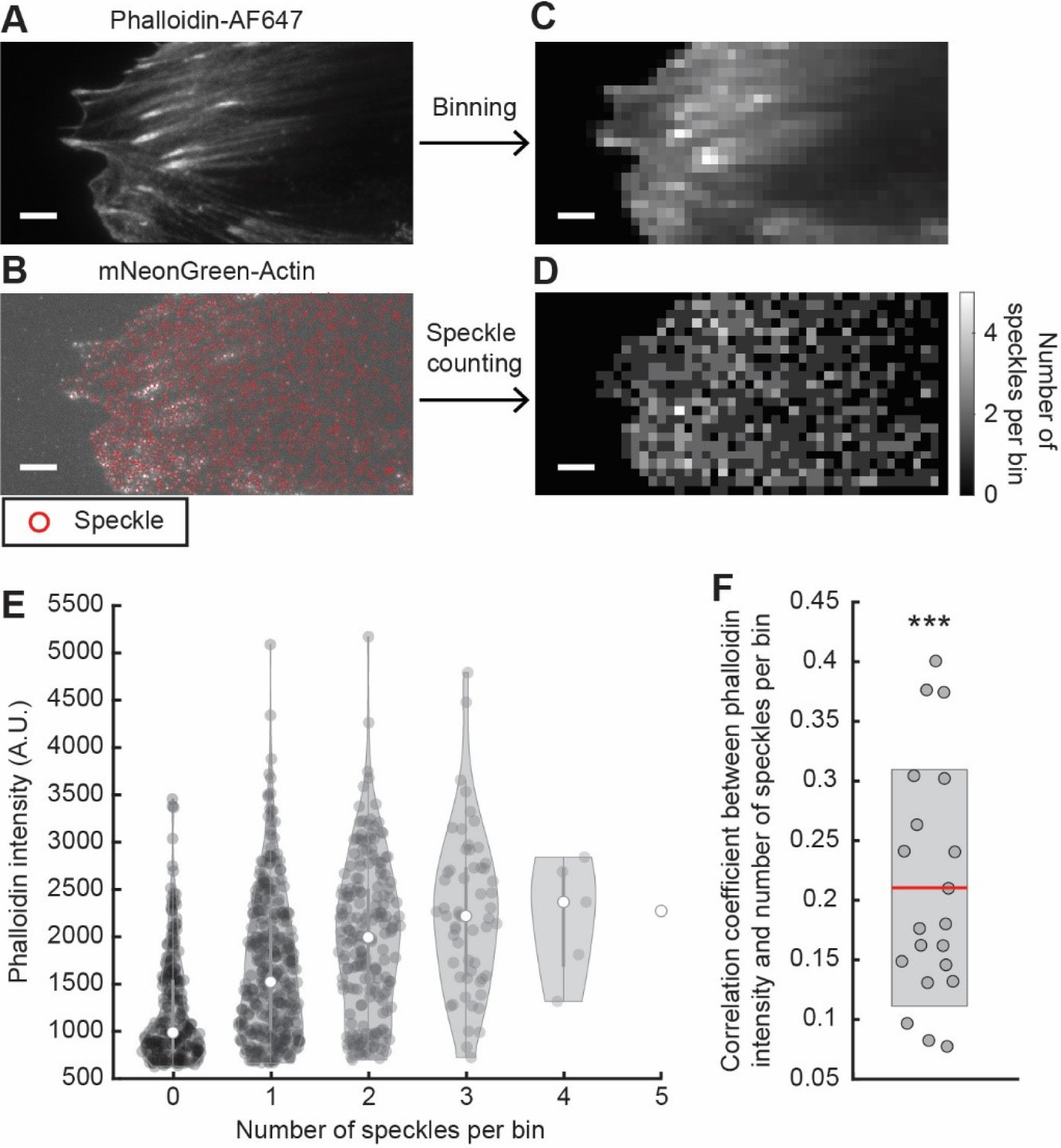
Correlation analysis of actin speckle density vs. CA density. **(A, B)** Representative images of Phalloidin-AF647 labelled actin **(A)** and actin speckles **(B)** in a TIME-mNGrActin cell. Red circles in B indicate detected speckles. Scale bar, 5 µm. **(C, D)** Both images were divided into ∼1.2 x 1.2 µm2 bins. Average phalloidin-AF647 intensity per bin is shown in **(C)** and number of speckles per bin is shown as a heatmap in **(D)**. Scale bar same as in A, B. **(E)** Violin plots of Phalloidin intensity per bin, in bins with indicated number of detected speckles, for a representative cell. Gray circles show individual bin measurements. White circles show median of group. **(F)** Correlation coefficient between phalloidin intensity and number of speckles per

**Figure S4.**
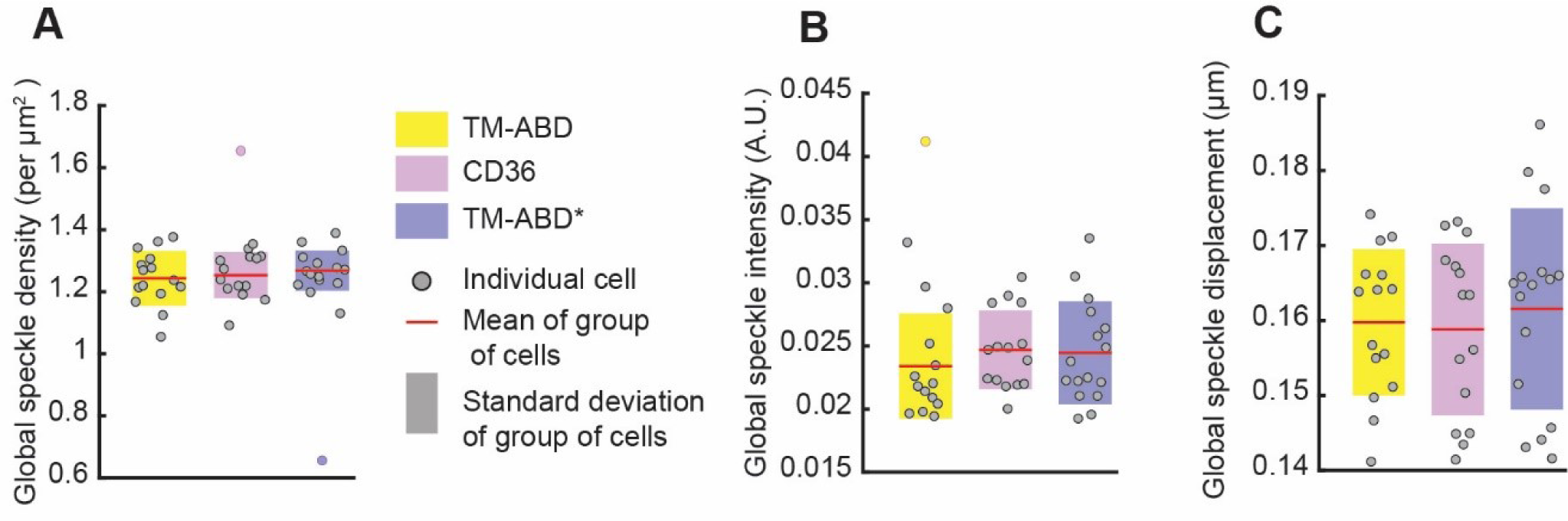
Global speckle properties in cells expressing CD36, TM-ABD or TM-ABD*. Global speckle **(A)** density, **(B)** intensity and **(C)** displacement magnitude of all actin speckles within the cell mask. Colors, circles, red lines, shaded bars and statistical tests as in Figs. 2 and 3. All p-values > 0.05 and are not explicitly indicated. See Table S8 for sample size and other dataset details.

**Figure S5.**
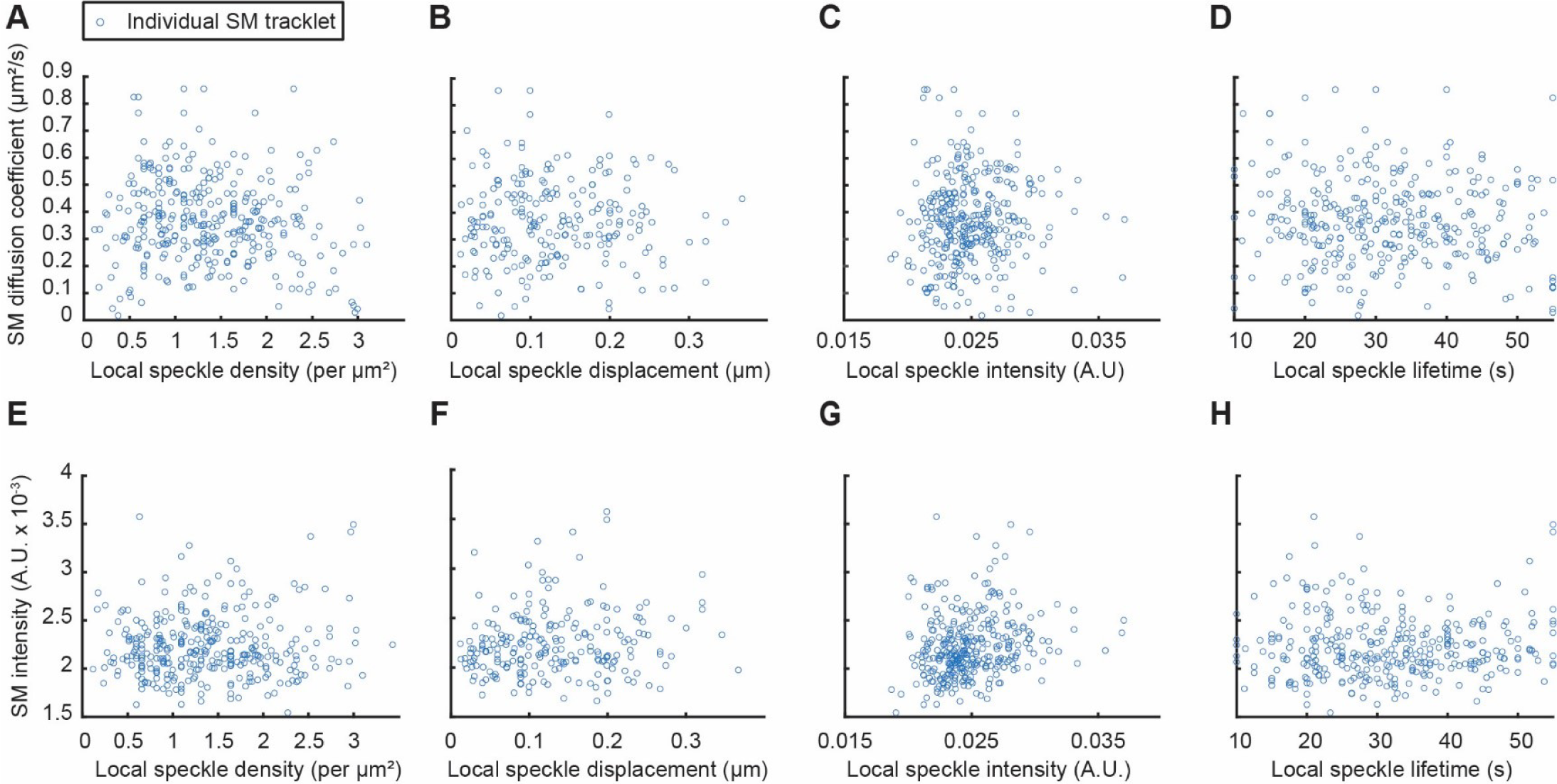
Scatterplots of indicated SM properties vs. associated local speckle properties from a representative single cell. **(A-D)** Scatterplots of SM tracklet diffusion coefficient vs local speckle density **(A)**, displacement magnitude **(B)**, intensity **(C)** and lifetime **(D)**. **(E-H)** Scatterplots of SM tracklet intensity vs local speckle density **(E)**, displacement magnitude **(F)**, intensity **(G)** and lifetime **(H)**. Each circle represents one free SM tracklet.

**Figure S6.**
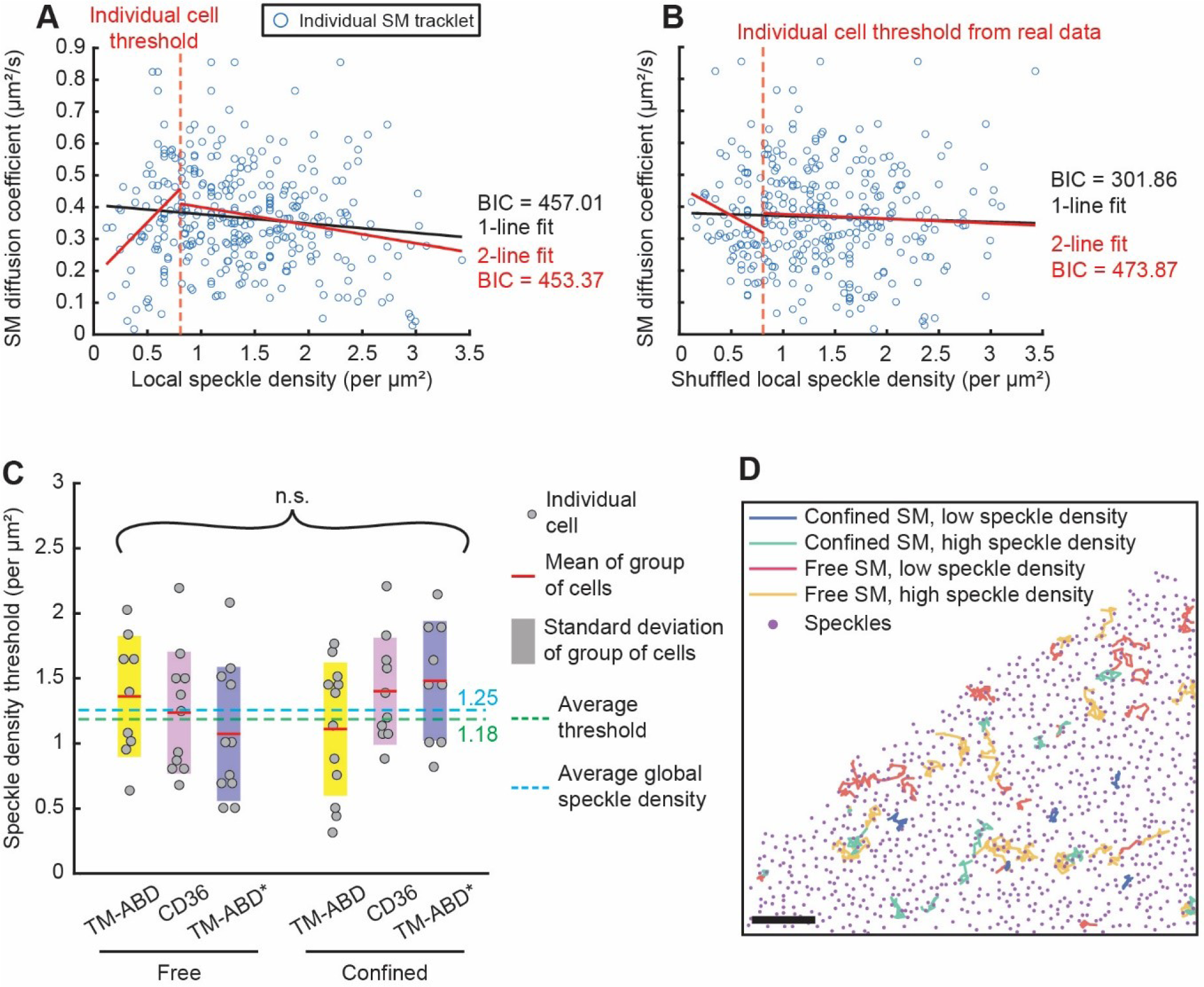
Assessment of biphasic behavior between SM diffusion coefficient and local speckle density. **(A, B)** Scatterplot of SM diffusion coefficient vs. local speckle density **(A)** or vs. shuffled local speckle density **(B)** from a single cell (same as in Fig. S5) with 1-line and 2-line fits. Each circle represents one free SM tracklet. Solid lines represent 1-line and 2-line fits with their BIC values, in black and red, respectively. Dashed vertical line indicates the local speckle density threshold derived from the 2-line fit for this cell (Suppl. Note 2). **(C)** Speckle density threshold per cell per PM protein type and diffusion type. Colors, circles, red lines and shaded bars as in Figs. 2 and 3. Comparison between PM protein types and diffusion types using ANOVA revealed no differences between them, with n.s. indicating p-value > 0.05. Green and cyan dashed lines indicate, respectively, the mean speckle density threshold and the mean global speckle density over all cells across the indicated groups. **(D)** CD36 tracklets in one FSM interval overlaid on speckle positions (at the beginning of FSM interval), colored by diffusion type and speckle density level. Scale bar: 5 µm. See Table S8 for sample size and other dataset details. See Suppl. Note 2 for biphasic analysis details.

**Figure S7.**
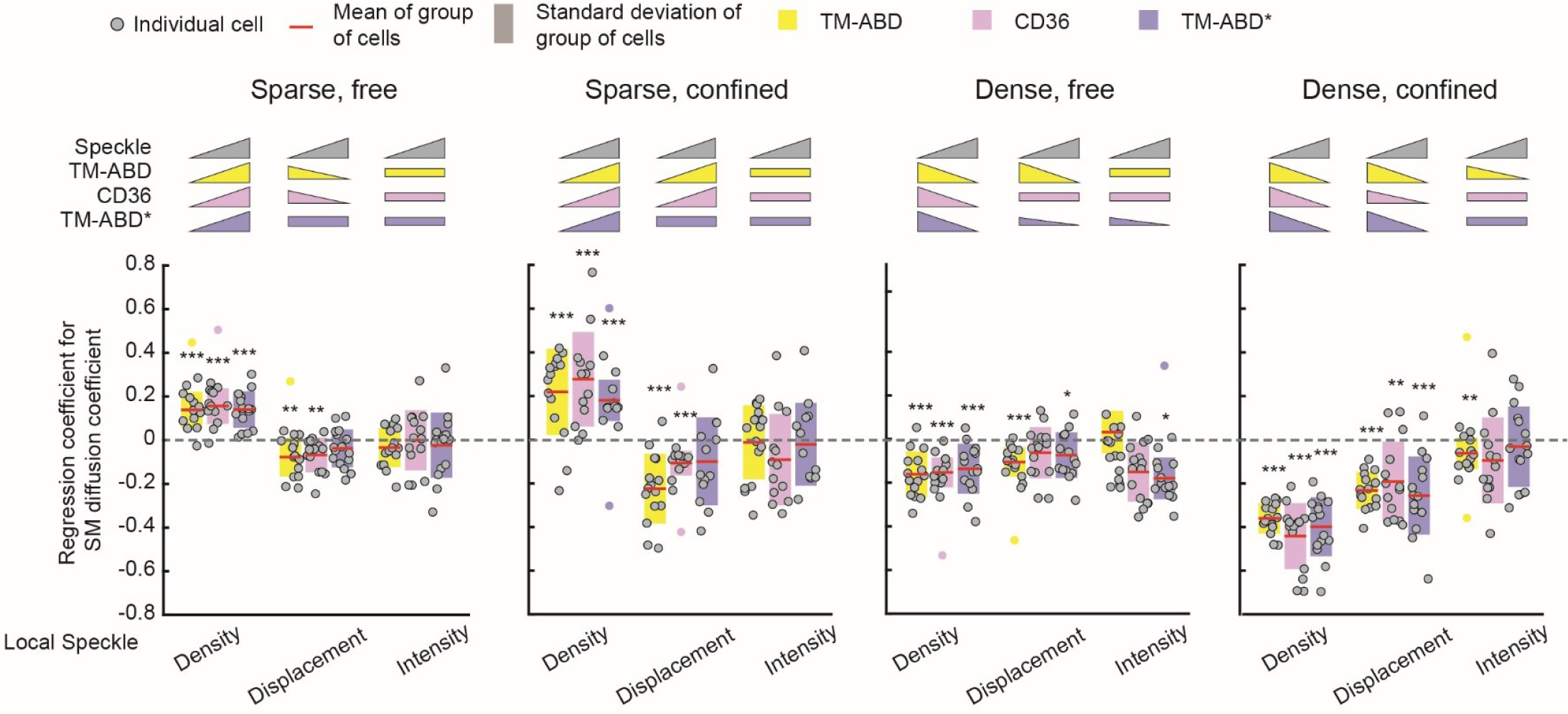
MVRG of SM diffusion coefficient on local speckle density, displacement magnitude and intensity. Coefficients from MVRG of SM diffusion coefficient on local speckle density, displacement magnitude and intensity for CD36 (magenta), TM-ABD (yellow) and TM-ABD* (blue). MVRG analysis was performed separately for the indicated subgroups of SM tracklets, based on diffusion type and local speckle density pairings. All details as in Fig. 4. SM diffusion coefficient showed very little dependence on speckle intensity. Inclusion of speckle intensity in MVRG analysis had little influence on the MVRG coefficients for speckle density and displacement magnitude (compare to Fig. 4A). See Table S8 for sample size and other dataset details.

**Figure S8.**
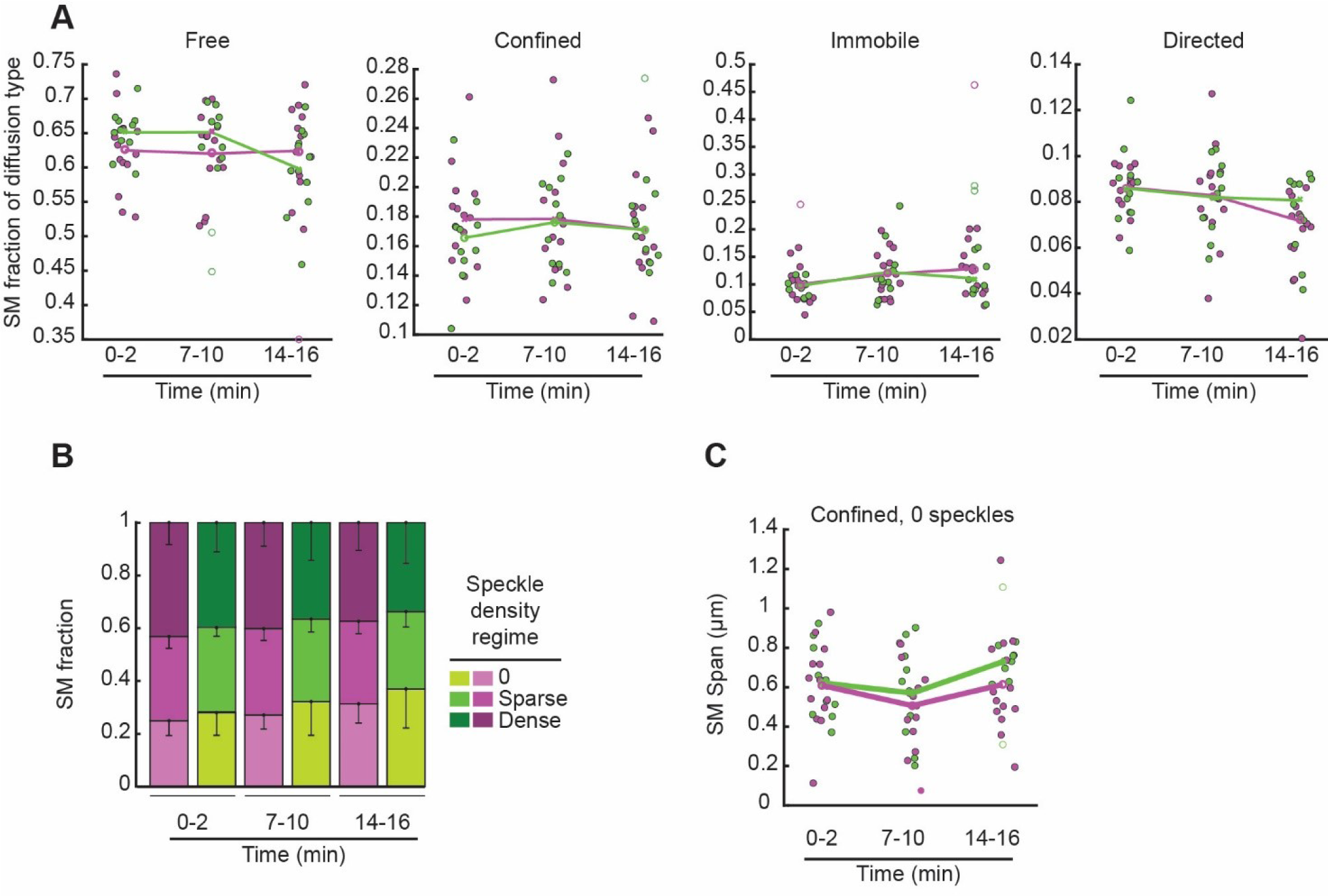
Further characterization of CD36 mobility in unperturbed and Latrunculin A perturbed cells. **(A)** Fraction of SM tracklets exhibiting the indicated diffusion types in the absence (magenta) or presence (green) of Lat A, at indicated instances. **(B)** Fraction of SM tracklets at different local speckle density levels. Each stacked bar represents the mean SM tracklet fraction over all cells for that condition and instance. Error bar, corresponding standard deviation. **(C)** Span of confined SM tracklets matched to 0 speckles. In A and C,

**Figure S9.**
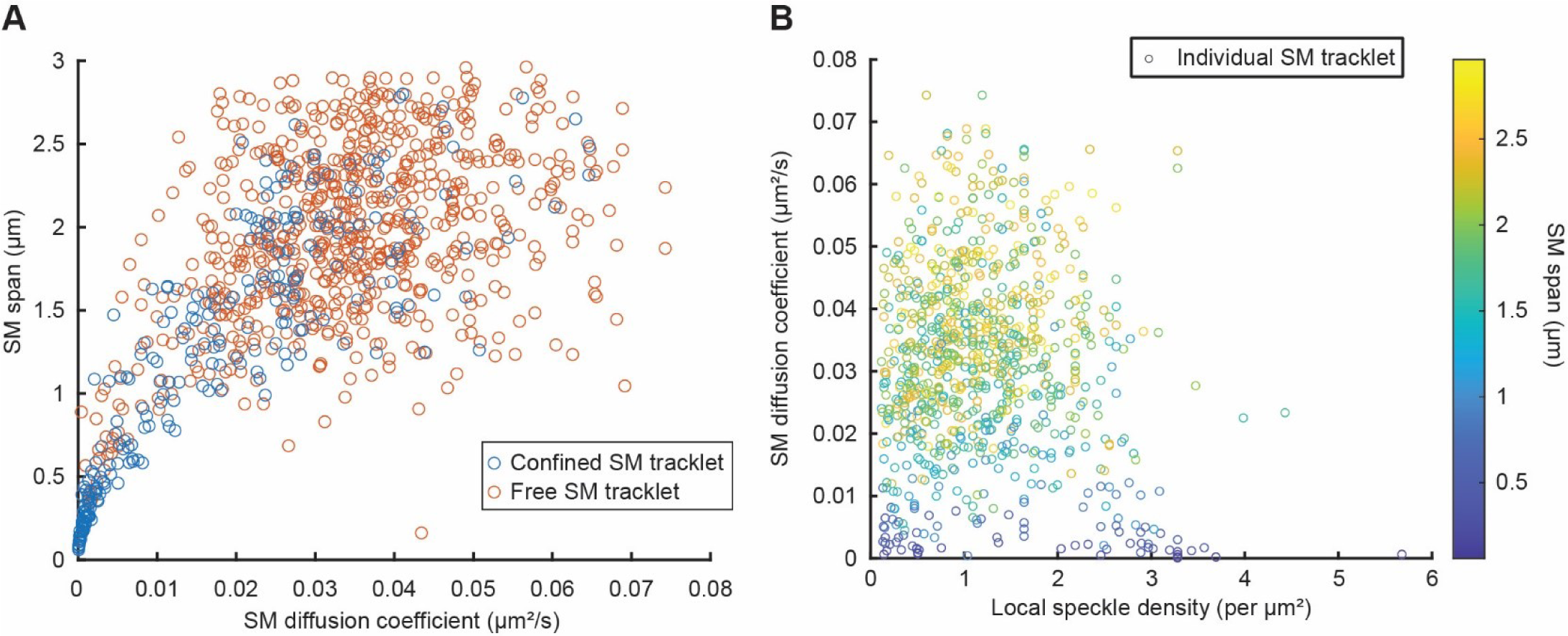
Relationship between SM diffusion coefficient, SM span and local speckle density. **(A)** Scatterplot of SM span vs SM diffusion coefficient for confined (blue) and free (red) SM tracklets from a representative cell. **(B)** Scatterplot of SM diffusion coefficient (free and confined SM tracklets combined) vs local speckle density colored by SM span from a representative cell.

## Supplemental tables

**Table S1.**
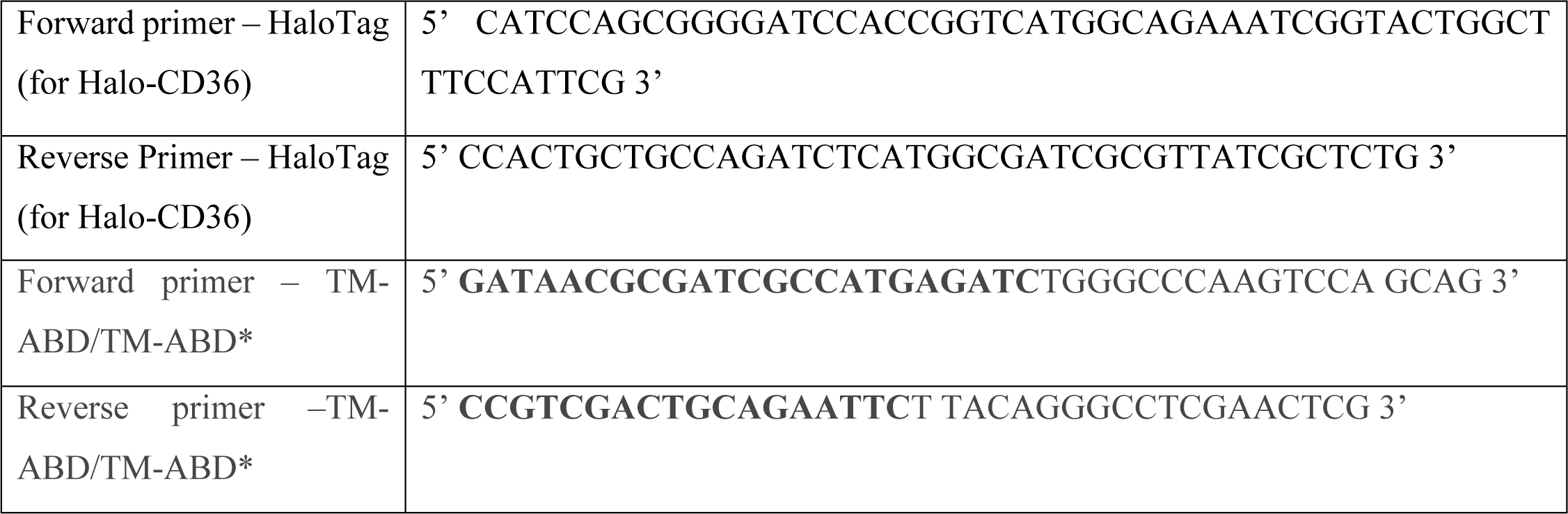
Primers used for infusion cloning.

**Table S2.** SM detection (u-track): Non-default parameter values.

|  |  |
| --- | --- |
| Gaussian standard deviation | 1.228 pixels (0.109 $\mu\text{m}$) |
| Gaussian mixture-model fitting at local maxima: Do iterative Gaussian mixture-model fitting | Yes |
| Gaussian mixture-model fitting at local maxima: α-value for residuals test | 0.01 |
| Gaussian mixture-model fitting at local maxima: α-value for distance test | 0.01 |

**Table S3.** SM tracking (u-track): Non-default parameter values.

|  |  |
| --- | --- |
| Overall: Do segment merging | Yes |
| Overall: Do segment splitting | Yes |
| Frame-to-frame linking: Brownian search radius upper and lower bound | 12 pixels (1.086 $\mu\text{m}$) |
| Gap closing, merging and splitting – general: Brownian search radius upper and lower bound | 12 pixels (1.068 $\mu\text{m}$) |
| Gap closing, merging and splitting – general: Scaling power to expand Brownian search radius | 0.1 |
| Gap closing, merging and splitting – general: Use nearest neighbor distance to expand Brownian search radius | No |
| Gap closing, merging and splitting – merging & splitting: Search radius lower bound | 3 pixels (0.267 $\mu\text{m}$) |
| Gap closing, merging and splitting – merging & splitting: For an end/start, the possibility of merging/splitting is allowed only if there is no possibility of gap closing | Yes<sup>a</sup> |
| Gap closing, merging and splitting – birth & death: Birth and death cost | Derived from gap closing and merging/splitting cost formulae |

<sup>a</sup> Limiting merging and splitting events to segment ends and starts that do not have the possibility of gap closing helps with distinguishing between molecular interactions and particles passing by each other.

**Table S4.** Speckle detection: Non-default parameter values.

|  |  |
| --- | --- |
| Standard deviation for Gaussian low-pass filtering | 0.096 $\mu\text{m}$ |
| Alpha value for statistical selection of speckles | 0.001 |
| Iterative speckle detection- Maximal number of iterations | 2 |
| Iterative speckle detection-Minimum fraction of speckles to be detected at each iteration | 0.01 |

**Table S5.**
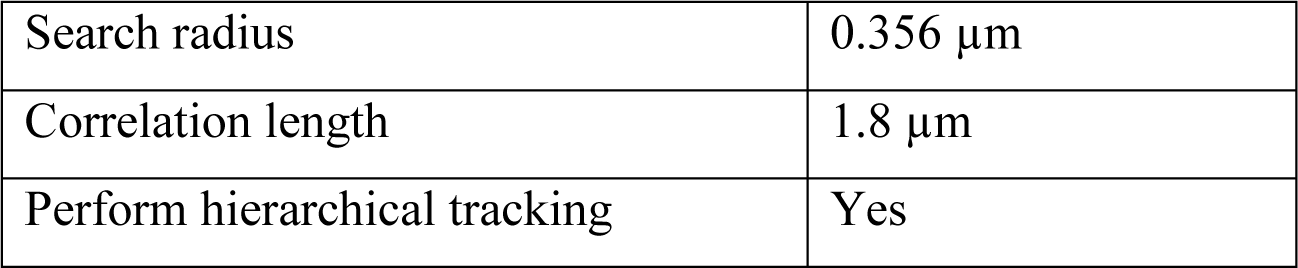
Speckle tracking: Non-default parameter values.

|  |  |
| --- | --- |
| Search radius | 0.356 $\mu\text{m}$ |
| Correlation length | 1.8 $\mu\text{m}$ |
| Perform hierarchical tracking | Yes |

**Table S6.** Speckle kinetic analysis: Non-default parameter values.

|  |  |
| --- | --- |
| Standard deviation for Gaussian low-pass filtering | 0.445 $\mu\text{m}$ |
| Number of frames for time averaging | 11 |

**Table S7.** Fraction of shuffled data MVRG coefficients with a p-value smaller than that of the real data MVRG coefficient. Black/gray entries indicate significant/non-significant MVRG coefficients.

| Figure | PM protein | Motion type | Speckle density type | PM protein SM property | Local speckle property | p-value | Fraction of p-values from 100 data shuffles that are smaller than real data p-value |
| --- | --- | --- | --- | --- | --- | --- | --- |
| 4A | TM-ABD | Free | Sparse | SM diffusion coefficient | Density | 6.95E-05 | 0.00 |
| 4A | CD36 | Free | Sparse | SM diffusion coefficient | Density | 1.76E-05 | 0.00 |
| 4A | TM-ABD* | Free | Sparse | SM diffusion coefficient | Density | 1.50E-06 | 0.00 |
| 4A | TM-ABD | Free | Sparse | SM diffusion coefficient | Displacement | 1.14E-02 | 0.03 |
| 4A | CD36 | Free | Sparse | SM diffusion coefficient | Displacement | 2.13E-02 | 0.02 |
| 4A | TM-ABD* | Free | Sparse | SM diffusion coefficient | Displacement | 1.35E-01 | 0.14 |
| 4A | TM-ABD | Confined | Sparse | SM diffusion coefficient | Density | 5.58E-04 | 0.00 |
| 4A | CD36 | Confined | Sparse | SM diffusion coefficient | Density | 1.70E-03 | 0.00 |
| 4A | TM-ABD* | Confined | Sparse | SM diffusion coefficient | Density | 2.40E-02 | 0.02 |
| 4A | TM-ABD | Confined | Sparse | SM diffusion coefficient | Displacement | 1.23E-05 | 0.00 |
| 4A | CD36 | Confined | Sparse | SM diffusion coefficient | Displacement | 1.66E-05 | 0.00 |
| 4A | TM-ABD* | Confined | Sparse | SM diffusion coefficient | Displacement | 2.01E-01 | 0.23 |
| 4A | TM-ABD | Free | Dense | SM diffusion coefficient | Density | 3.20E-05 | 0.00 |
| 4A | CD36 | Free | Dense | SM diffusion coefficient | Density | 9.28E-07 | 0.00 |
| 4A | TM-ABD* | Free | Dense | SM diffusion coefficient | Density | 4.02E-04 | 0.00 |
| 4A | TM-ABD | Free | Dense | SM diffusion coefficient | Displacement | 1.26E-05 | 0.00 |
| 4A | CD36 | Free | Dense | SM diffusion coefficient | Displacement | 2.58E-02 | 0.06 |
| 4A | TM-ABD* | Free | Dense | SM diffusion coefficient | Displacement | 2.59E-02 | 0.02 |
| 4A | TM-ABD | Confined | Dense | SM diffusion coefficient | Density | 5.07E-11 | 0.00 |
| 4A | CD36 | Confined | Dense | SM diffusion coefficient | Density | 5.41E-08 | 0.00 |
| 4A | TM-ABD* | Confined | Dense | SM diffusion coefficient | Density | 1.34E-08 | 0.00 |
| 4A | TM-ABD | Confined | Dense | SM diffusion coefficient | Displacement | 1.61E-07 | 0.00 |
| 4A | CD36 | Confined | Dense | SM diffusion coefficient | Displacement | 7.04E-03 | 0.04 |
| 4A | TM-ABD* | Confined | Dense | SM diffusion coefficient | Displacement | 1.01E-04 | 0.00 |
| 4B | TM-ABD | Free | Sparse | SM intensity | Density | 8.03E-01 | 0.85 |
| 4B | CD36 | Free | Sparse | SM intensity | Density | 1.17E-01 | 0.17 |
| 4B | TM-ABD* | Free | Sparse | SM intensity | Density | 5.24E-01 | 0.59 |
| 4B | TM-ABD | Free | Sparse | SM intensity | Displacement | 2.19E-01 | 0.27 |
| 4B | CD36 | Free | Sparse | SM intensity | Displacement | 4.43E-01 | 0.46 |
| 4B | TM-ABD* | Free | Sparse | SM intensity | Displacement | 5.11E-01 | 0.55 |
| 4B | TM-ABD | Free | Sparse | SM intensity | Intensity | 2.64E-04 | 0.00 |
| 4B | CD36 | Free | Sparse | SM intensity | Intensity | 1.19E-04 | 0.00 |
| 4B | TM-ABD* | Free | Sparse | SM intensity | Intensity | 1.24E-05 | 0.00 |
| 4B | TM-ABD | Confined | Sparse | SM intensity | Density | 8.94E-02 | 0.19 |
| 4B | CD36 | Confined | Sparse | SM intensity | Density | 6.50E-02 | 0.12 |
| 4B | TM-ABD* | Confined | Sparse | SM intensity | Density | 4.56E-01 | 0.48 |
| 4B | TM-ABD | Confined | Sparse | SM intensity | Displacement | 3.24E-01 | 0.39 |
| 4B | CD36 | Confined | Sparse | SM intensity | Displacement | 2.56E-02 | 0.06 |
| 4B | TM-ABD* | Confined | Sparse | SM intensity | Displacement | 7.33E-01 | 0.76 |
| 4B | TM-ABD | Confined | Sparse | SM intensity | Intensity | 7.88E-03 | 0.02 |
| 4B | CD36 | Confined | Sparse | SM intensity | Intensity | 1.26E-05 | 0.00 |
| 4B | TM-ABD* | Confined | Sparse | SM intensity | Intensity | 7.36E-01 | 0.74 |
| 4B | TM-ABD | Free | Dense | SM intensity | Density | 2.42E-01 | 0.28 |
| 4B | CD36 | Free | Dense | SM intensity | Density | 2.83E-03 | 0.00 |
| 4B | TM-ABD* | Free | Dense | SM intensity | Density | 2.69E-01 | 0.29 |
| 4B | TM-ABD | Free | Dense | SM intensity | Displacement | 3.02E-03 | 0.00 |
| 4B | CD36 | Free | Dense | SM intensity | Displacement | 9.74E-02 | 0.09 |
| 4B | TM-ABD* | Free | Dense | SM intensity | Displacement | 6.45E-01 | 0.59 |
| 4B | TM-ABD | Free | Dense | SM intensity | Intensity | 3.29E-06 | 0.00 |
| 4B | CD36 | Free | Dense | SM intensity | Intensity | 3.34E-07 | 0.00 |
| 4B | TM-ABD* | Free | Dense | SM intensity | Intensity | 9.60E-07 | 0.00 |
| 4B | TM-ABD | Confined | Dense | SM intensity | Density | 8.36E-02 | 0.15 |
| 4B | CD36 | Confined | Dense | SM intensity | Density | 2.95E-03 | 0.01 |
| 4B | TM-ABD* | Confined | Dense | SM intensity | Density | 9.50E-01 | 0.94 |
| 4B | TM-ABD | Confined | Dense | SM intensity | Displacement | 8.24E-03 | 0.00 |
| 4B | CD36 | Confined | Dense | SM intensity | Displacement | 7.70E-01 | 0.88 |
| 4B | TM-ABD* | Confined | Dense | SM intensity | Displacement | 5.93E-01 | 0.61 |
| 4B | TM-ABD | Confined | Dense | SM intensity | Intensity | 1.53E-04 | 0.00 |
| 4B | CD36 | Confined | Dense | SM intensity | Intensity | 6.72E-03 | 0.01 |
| 4B | TM-ABD* | Confined | Dense | SM intensity | Intensity | 1.18E-01 | 0.11 |
| S7 | TM-ABD | Free | Sparse | SM diffusion coefficient | Density | 2.92E-05 | 0.00 |
| S7 | CD36 | Free | Sparse | SM diffusion coefficient | Density | 1.54E-05 | 0.00 |
| S7 | TM-ABD* | Free | Sparse | SM diffusion coefficient | Density | 6.83E-06 | 0.00 |
| S7 | TM-ABD | Free | Sparse | SM diffusion coefficient | Displacement | 6.24E-03 | 0.03 |
| S7 | CD36 | Free | Sparse | SM diffusion coefficient | Displacement | 6.58E-03 | 0.00 |
| S7 | TM-ABD* | Free | Sparse | SM diffusion coefficient | Displacement | 1.05E-01 | 0.13 |
| S7 | TM-ABD | Free | Sparse | SM diffusion coefficient | Intensity | 1.71E-01 | 0.23 |
| S7 | CD36 | Free | Sparse | SM diffusion coefficient | Intensity | 9.94E-01 | 0.99 |
| S7 | TM-ABD* | Free | Sparse | SM diffusion coefficient | Intensity | 5.48E-01 | 0.54 |
| S7 | TM-ABD | Confined | Sparse | SM diffusion coefficient | Density | 7.53E-04 | 0.00 |
| S7 | CD36 | Confined | Sparse | SM diffusion coefficient | Density | 5.87E-04 | 0.01 |
| S7 | TM-ABD* | Confined | Sparse | SM diffusion coefficient | Density | 8.55E-05 | 0.00 |
| S7 | TM-ABD | Confined | Sparse | SM diffusion coefficient | Displacement | 8.96E-05 | 0.00 |
| S7 | CD36 | Confined | Sparse | SM diffusion coefficient | Displacement | 9.20E-05 | 0.00 |
| S7 | TM-ABD* | Confined | Sparse | SM diffusion coefficient | Displacement | 1.00E-01 | 0.05 |
| S7 | TM-ABD | Confined | Sparse | SM diffusion coefficient | Intensity | 7.95E-01 | 0.82 |
| S7 | CD36 | Confined | Sparse | SM diffusion coefficient | Intensity | 1.42E-01 | 0.25 |
| S7 | TM-ABD* | Confined | Sparse | SM diffusion coefficient | Intensity | 6.91E-01 | 0.77 |
| S7 | TM-ABD | Free | Dense | SM diffusion coefficient | Density | 3.45E-05 | 0.00 |
| S7 | CD36 | Free | Dense | SM diffusion coefficient | Density | 1.23E-06 | 0.00 |
| S7 | TM-ABD* | Free | Dense | SM diffusion coefficient | Density | 4.08E-04 | 0.00 |
| S7 | TM-ABD | Free | Dense | SM diffusion coefficient | Displacement | 5.58E-05 | 0.00 |
| S7 | CD36 | Free | Dense | SM diffusion coefficient | Displacement | 5.68E-02 | 0.04 |
| S7 | TM-ABD* | Free | Dense | SM diffusion coefficient | Displacement | 1.61E-02 | 0.01 |
| S7 | TM-ABD | Free | Dense | SM diffusion coefficient | Intensity | 2.15E-01 | 0.25 |
| S7 | CD36 | Free | Dense | SM diffusion coefficient | Intensity | 4.66E-01 | 0.46 |
| S7 | TM-ABD* | Free | Dense | SM diffusion coefficient | Intensity | 4.68E-02 | 0.07 |
| S7 | TM-ABD | Confined | Dense | SM diffusion coefficient | Density | 1.04E-11 | 0.00 |
| S7 | CD36 | Confined | Dense | SM diffusion coefficient | Density | 6.12E-08 | 0.00 |
| S7 | TM-ABD* | Confined | Dense | SM diffusion coefficient | Density | 1.75E-08 | 0.00 |
| S7 | TM-ABD | Confined | Dense | SM diffusion coefficient | Displacement | 4.82E-08 | 0.00 |
| S7 | CD36 | Confined | Dense | SM diffusion coefficient | Displacement | 1.69E-03 | 0.00 |
| S7 | TM-ABD* | Confined | Dense | SM diffusion coefficient | Displacement | 7.16E-05 | 0.00 |
| S7 | TM-ABD | Confined | Dense | SM diffusion coefficient | Intensity | 9.03E-03 | 0.01 |
| S7 | CD36 | Confined | Dense | SM diffusion coefficient | Intensity | 9.22E-02 | 0.09 |
| S7 | TM-ABD* | Confined | Dense | SM diffusion coefficient | Intensity | 5.21E-01 | 0.51 |

**Table S8.**
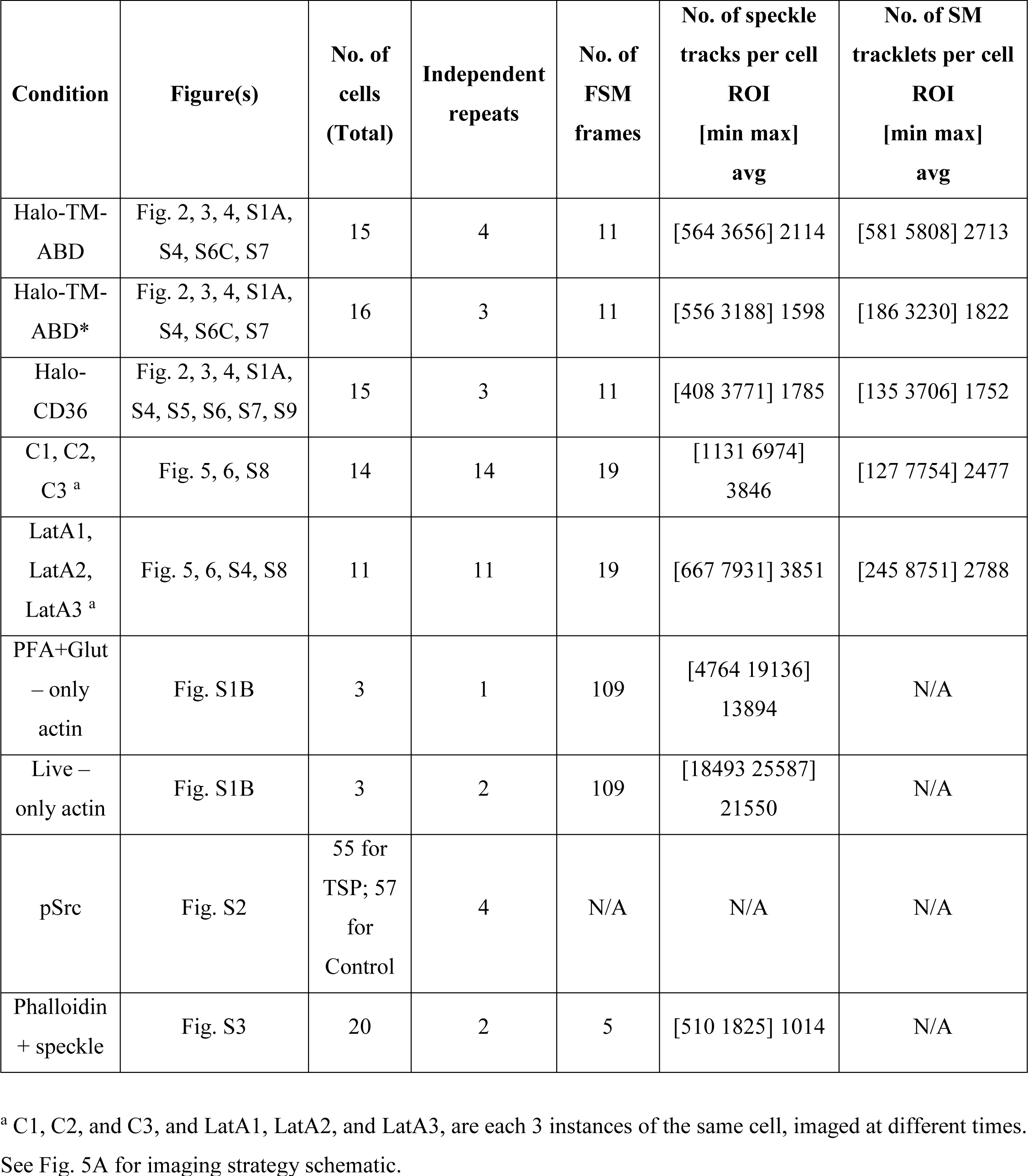
Dataset properties for the different experimental conditions.

| **Condition** | **Figure(s)** | **No. of cells (Total)** | **Independent repeats** | **No. of FSM frames** | **No. of speckle tracks per cell ROI [min max] avg** | **No. of SM tracklets per cell ROI [min max] avg** |
| --- | --- | --- | --- | --- | --- | --- |
| Halo-TM-ABD | Fig. 2, 3, 4, S1A, S4, S6C, S7 | 15 | 4 | 11 | [564 3656] 2114 | [581 5808] 2713 |
| Halo-TM-ABD* | Fig. 2, 3, 4, S1A, S4, S6C, S7 | 16 | 3 | 11 | [556 3188] 1598 | [186 3230] 1822 |
| Halo-CD36 | Fig. 2, 3, 4, S1A, S4, S5, S6, S7, S9 | 15 | 3 | 11 | [408 3771] 1785 | [135 3706] 1752 |
| C1, C2, C3 <sup>a</sup> | Fig. 5, 6, S8 | 14 | 14 | 19 | [1131 6974] 3846 | [127 7754] 2477 |
| LatA1, LatA2, LatA3 <sup>a</sup> | Fig. 5, 6, S4, S8 | 11 | 11 | 19 | [667 7931] 3851 | [245 8751] 2788 |
| PFA+Glut – only actin | Fig. S1B | 3 | 1 | 109 | [4764 19136] 13894 | N/A |
| Live – only actin | Fig. S1B | 3 | 2 | 109 | [18493 25587] 21550 | N/A |
| pSrc | Fig. S2 | 55 for TSP; 57 for Control | 4 | N/A | N/A | N/A |
| Phalloidin + speckle | Fig. S3 | 20 | 2 | 5 | [510 1825] 1014 | N/A |

<sup>a</sup> C1, C2, and C3, and LatA1, LatA2, and LatA3, are each 3 instances of the same cell, imaged at different times. See Fig. 5A for imaging strategy schematic.

